# A Transparent and Generalizable Deep Learning Framework for Genomic Ancestry Prediction

**DOI:** 10.1101/2025.08.26.672448

**Authors:** Camille Rochefort-Boulanger, Matthew Scicluna, Raphaël Poujol, Jean-Christophe Grenier, Pierre Luc Carrier, Sébastien Lemieux, Julie G Hussin

**Affiliations:** Research Centre, Montreal Heart Institute, Montreal, QC, Canada; Département de Biochimie et Médecine Moléculaire, Université de Montréal, Montreal, QC, Canada; Mila - Quebec Artificial Intelligence Institute, Montreal, QC, Canada; Institute for Research in Immunology and Cancer, Université de Montréal, Montreal, QC, Canada; Département de Médecine, Université de Montréal, Montreal, QC, Canada

## Abstract

Accurately capturing genetic ancestry is critical for ensuring reproducibility and fairness in genomic studies and downstream health research. This study aims to address the prediction of ancestry from genetic data using deep learning, with a focus on generalizability across datasets with diverse populations and on explainability to improve model transparency. We adapt the Diet Network, a deep learning architecture proven effective in handling high-dimensional data, to learn population ancestry from single nucleotide polymorphisms (SNPs) data using the populational Thousand Genomes Project dataset. Our results highlight the model’s ability to generalize to diverse populations in the CARTaGENE and Montreal Heart Institute biobanks and that predictions remain robust to high levels of missing SNPs. We show that, despite the lack of North African populations in the training dataset, the model learns latent representations that reflect meaningful population structure for North African individuals in the biobanks. To improve model transparency, we apply Saliency Maps, DeepLift, GradientShap and Integrated Gradients attribution techniques and evaluate their performance in identifying SNPs leveraged by the model. Using DeepLift, we show that model’s predictions are driven by population-specific signals consistent with those identified by traditional population genetics metrics. This work presents a generalizable and interpretable deep learning framework for genetic ancestry inference in large-scale biobanks with genetic data. By enabling more widespread genomic ancestry characterization in these cohorts, this study contributes practical tools for integrating genetic data into downstream biomedical applications, supporting more inclusive and equitable healthcare solutions.

## 2 Introduction

With the rapid advancement of high-throughput sequencing and genotyping technologies [1, 2], large-scale clinical cohorts and biobank datasets containing genetic data have become increasingly available, opening new opportunities for large-scale, data-driven health research [3]. These resources have underscored the importance of including individuals from diverse genetic ancestry groups to ensure the validity, reproducibility, and clinical relevance of genetic findings [4, 5]. In response, multiple initiatives have emerged to build biobanks that better represent populations historically underrepresented in genomic research [6, 7, 8]. These efforts have also spurred interest in accurately characterizing population structure and inferring genetic ancestry, especially since population labels in biobanks are often based on self-reported ethnicity [9]. Since such labels can be incomplete, imprecise, or constrained by predefined categories in questionnaires, genetic ancestry inference from genomic data offers a complementary, biologically meaningful, and often more consistent alternative [10].

Population labels have long been used in genetics to define reference populations, providing a framework for comparing genetic diversity across groups [11, 12, 13]. These labels guide the study of population structure [14, 15], evolutionary history [16, 17], and disease susceptibility [18, 19, 20]. Publically available population reference datasets, such as HapMap [21], the Thousand Genomes Project (1KGP) [22] and the Human Genome Diversity Project (HGDP) [23], play a central role in human genetics by offering well-curated genetic data from individuals of diverse geographic and ancestral backgrounds. In these reference datasets, population labels are typically defined using a combination of geographic origin, linguistic background, and self-reported ancestry.

Traditional methods which often rely on statistical techniques like principal component analysis (PCA) [24, 25] and local ancestry inference tools [26], depend on these reference panels with population labels. More recently, the rise of data-hungry deep learning methods applied to such data is also enabling a new generation of genomics research [27, 28, 29] including efforts to infer genetic ancestry [30, 31]. However, the application of traditional and deep learning methods for inferring genetic ancestry in biobanks presents limitations. These inferences have limited generalizability across datasets, due to the reliance of those methods on fixed sets of SNPs that will inevitably differ between cohorts, requiring reselecting SNPs in a reference panel, followed by retraining.

Deep learning models are also prone to overfitting due to the “fat data problem”, where the number of genetic variants, such as single nucleotide polymorphisms (SNPs), far exceeds the number of samples [28]. Furthermore, deep learning models are often criticized as “black boxes” due to their limited interpretability, providing little transparency into the specific SNPs driving their predictions [32].

Another challenge in genetic ancestry inference is the reliance on discrete population labels. While these labels enable group-level comparisons, they fail to capture the continuum of human genetic variation, particularly for individuals from admixed or underrepresented populations. This oversimplification may introduce biases in downstream analyses, including health-related predic-tions, and carries the risk of reinforcing discriminatory interpretations.

In this study, we address the challenges of generalizability and interpretability in genetic ancestry inference by adapting the Diet Network [28], a deep learning framework designed to handle high-dimensional genetic data. The Diet Network reduces overfitting by leveraging per-SNP summary statistics to constrain the network’s weights, enabling efficient training even when the number of SNPs far exceeds the number of samples. We evaluated its generalizability across a diverse spectrum of populations included in two biobanks, the Montreal Heart Institute Biobank (MHIB) [33, 34] and the CARTaGENE (CaG) biobank [35, 36], which reflect the genetic heterogeneity of clinical cohorts. We demonstrated that the models maintain consistent predictions when applied to reduced sets of SNPs, highlighting their robustness to differences in genetic marker coverage across datasets. Additionally, we introduce a visualization method that captures how the model represents genetic diversity, an approach that uses the structure learned from the data to move beyond rigid population labels, extending the approach to individuals from admixed populations not represented in the training dataset. To improve transparency, we apply local feature attribution methods to identify SNPs that are most predictive of population labels. By comparing these attributions with traditional population genetics metrics, we gain a deeper understanding of how deep learning models capture population structure. This framework aims to address the technical challenges associated with both the use of genetic data with explainable deep learning and the issues with population labelling in current datasets, contributing to more transparent and generalizable predictions in genomics.

## 3 Data Description

We used genetic data from three distinct sources, each offering complementary strengths in terms of diversity, resolution, and real-world applicability. The 1KGP dataset was used to train our models and the generalizability of these trained models was evaluated in two large-scale Canadian cohorts: CaG and MHIB. Each dataset includes genetic data of participants obtained through genotyping arrays or whole genome sequencing, as well as self-reported information on ethnicity.

### 3.1 The 1KGP Populational Dataset

The 1KGP populational dataset [22] is a publicly available resource of human genetic variation providing high-resolution data from diverse populations and widely used to describe human genetic diversity. The dataset includes samples from 26 populations, grouped into superpopulations representing five continental ancestry groups (Europe, East Asia, South Asia, Africa and America). Participants were included if they self-reported having at least three out of four grand-parents from the same population. Population descriptions along with sample inclusion criteria and the guidelines for referring to these populations are available on the NHGRI website (https://www.coriell.org/1/NHGRI). Supplementary Table 1 lists the population descriptors and their corresponding three-letter abbreviation code.

1KGP provides genotyping data of 3,450 samples and high-coverage whole genome sequencing of 2,504 samples, with partial overlap between the two groups [37]. The Diet Network was previously trained [28] using genotyping data from 3,450 samples to maximize sample size for model training. During data review, we identified a small number of genotyping inconsistencies that had passed standard quality control filters (Supplementary Figure 1) and which were also reported in [38]. As a result, we focused on high coverage whole genome sequencing data in this study, prioritizing data quality over sample quantity. This data is available freely for download through the International Genome Sample Resource website (https://www.internationalgenome.org).

### 3.2 Canadian Biobanks

CaG and MHIB are two major Canadian research resources based in the province of Quebec. Both are managed by institutions located in Quebec and primarily include participants of French Canadian ancestry, a population of European descent. In addition to this majority group, both biobanks also include individuals from diverse genetic backgrounds reflecting the broader demographic landscape of the province.

The CaG biobank provides comprehensive data on environmental, lifestyle and genetic factors to support chronic disease research. Participants self-reported their ethnicity in predefined categories as well as information on the country of birth of their parents and grand-parents (Supplementary Methods **S1.1**). While the CaG platform includes genotype data for 29,334 individuals, these were obtained using different genotyping arrays over time. For this study, we focused on the subset of 17,286 individuals genotyped using the same Illumina Infinium Global Screening Array (GSA), in order to ensure consistency in variant coverage across individuals. Researchers can request access to the data by submitting a project proposal through the study website (www.cartagene.qc.ca). The MHIB, established to advance research on cardiovascular disease and genetics, primarily consists of hospital patients, along with individuals recruited from the general population. Participants self-reported their ethnicity in predefined categories (Supplementary Method **S1.1**) and indicated whether they are of French Canadian descent, specifying whether one or both parents are of French Canadian descent. Genotype data for 16,707 samples generated using the same Illumina Infinium GSA array version as the selected CaG subset are available through a formal project submission process in collaboration with a researcher affiliated with the Montreal Heart Institute. This requirement ensures compliance with the ethical consent provided by participants and the governance framework established for the use of biobank data (https://icm-mhi.org). Researchers interested in accessing the data are encouraged to contact the corresponding author for guidance on the access process.

## 4 Results

### 4.1 Merging Labels of Genetically Similar Populations

The 1KGP dataset includes genetically similar populations with distinct labels that reflect geographic separation rather than genetic differences. Training models to distinguish between such labels can lead to overfitting, where models rely on subtle allele frequency differences rather than capturing robust population structure, ultimately reducing the model’s ability to generalize to new data. To address this issue, we redefined some of the population labels by merging genetically similar populations under a single label. Specifically, we combined the European CEU and GBR populations into a single label (CEUGBR) and the South Asian STU and ITU populations into a single label (STUITU) using a combination of quantitative and qualitative criteria: low pairwise genetic differentiation using *F*_*ST*_ [39, 40], PCA clustering, and evidence of close familial relationships (Supplementary Results **S2.1** and Supplementary Figure 2).

### 4.2 Training Models Robust to Missing Data

Diet Network (Supplementary Figure 3) models were trained on 1KGP to classify 2,322 individuals into one of 24 populations using 229,986 SNPs. Training was performed using 5-fold cross-validation and three repetitions were analyzed, resulting in 15 models (Supplementary Methods **S1.2**). To improve the robustness of the models to missing genotypes, whether due to technical artifacts or non-overlapping SNPs across datasets, we trained the models using input dropout [41]. This regularization technique randomly masks input feature information (genotypes) during training to prevent the model from relying too heavily on a small subset of informative SNPs. Models were trained with varying input dropout rates, and their ability to handle missing SNPs was assessed by measuring classification accuracy on a held-out test set, in which missing values were simulated by randomly removing increasing proportions of SNPs (Supplementary Methods **S1.2**).

Models trained with input dropout demonstrated improved performances and a greater robustness to missing values (Figure 1A). More precisely, an input dropout rate of 0.995, retaining an average of 1,150 SNPs per sample during training, yielded the best accuracy on the complete test set (see Supplementary Figure 4 for models’ predictions) and maintained higher accuracy even with up to 99% simulated missing SNPs. Increasing the input dropout rate further led to a decline in performance.

**Figure 1:**
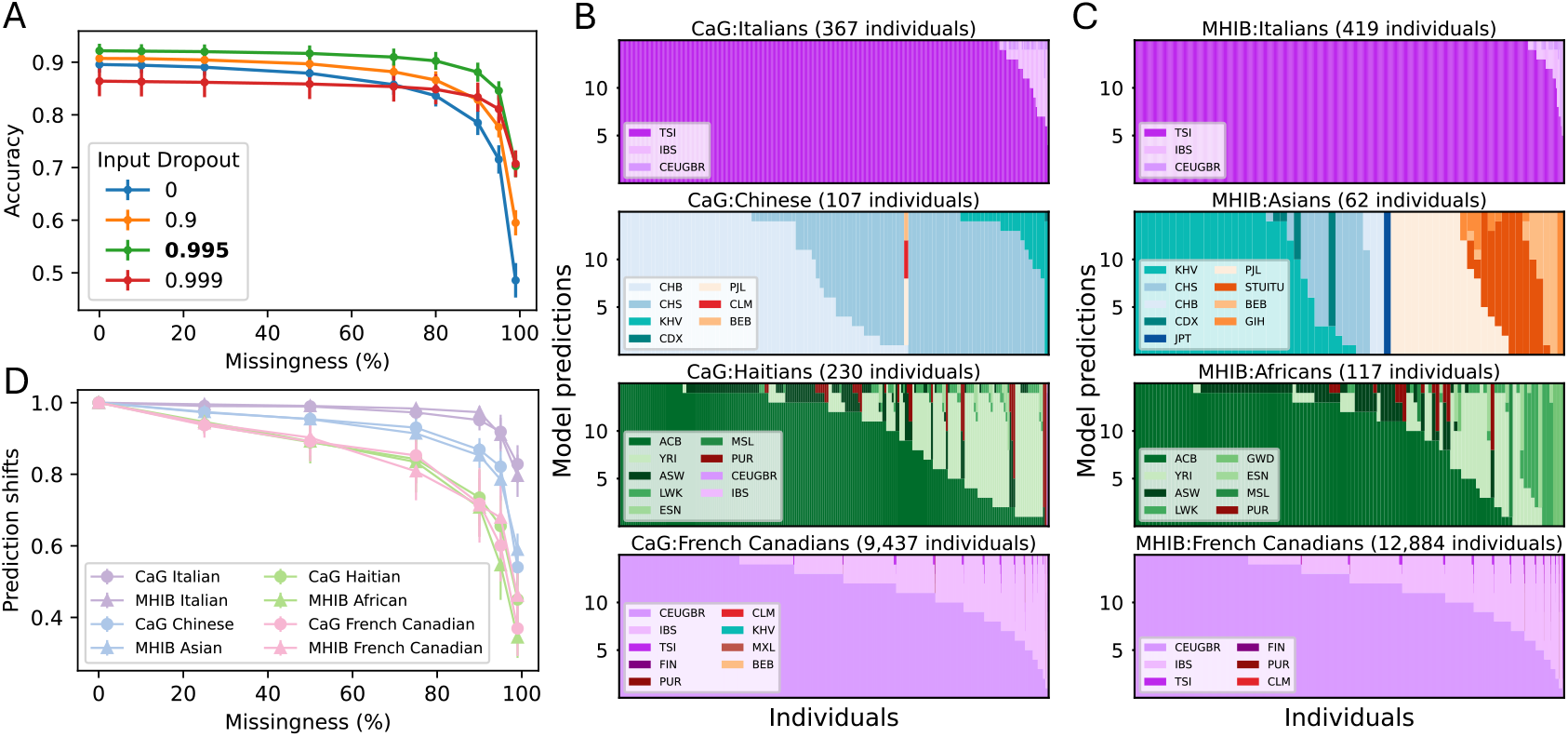
Diet Network Generalization. **(A)** Classification accuracy (y-axis) of the Diet Network models under increasing proportions of missing SNPs (x-axis), evaluated for input dropout rates 0, 0.9, 0.995 and 0.999. Accuracies at 0% missingness are averaged over 15 models, while accuracies for non-zero missingness proportions are averaged over the same 15 models and 100 distinct random SNP removal sets. **(B-C)** Predicted population labels for CaG (B) and MHIB (C) individuals. Each vertical line represents an individual and the y-axis shows the population predicted by each of the 15 models. **(D)** Proportion of models maintaining the same prediction as as with the complete SNP data (y-axis) across increasing levels of missing SNPs (x-axis).

### 4.3 Validating Generalizability in External Datasets

We assessed the generalizability of Diet Network models trained on the 1KGP dataset by applying the best-performing models (trained with 99.5% input dropout) to individuals with diverse self-reported ancestries from the CaG and MHIB datasets (Supplementary Methods **S1.1** and **S1.2**).

All 229, 986 SNPs used for training were selected to be shared across the three datasets to ensure a consistent evaluation (see Methods).

#### 4.3.1 Model predictions capture continental ancestry

##### Italian population

For individuals of Italian ancestry, present in both CaG and MHIB, model predictions were consistently assigned to the corresponding Italian TSI population in 1KGP, with some models occasionally assigning nearby European populations (Figure 1B–C, Italians).

##### Chinese and Asian populations

Among individuals of Chinese ancestry in CaG, the predicted populations aligned with East Asian populations from 1KGP such as CHB and CHS (Figure 1B, CaG:Chinese). In MHIB, the broad self-reported Asian category encompassed both East and South Asian ancestries, reflected in the model predictions which spanned EAS and SAS superpopulations (Figure 1C, MHIB: Asian). Across MHIB Asians individuals, all fifteen models consistently predicted only EAS or SAS populations indicating agreement between models at the continental level.

##### Haitian and African populations

Predictions of Haitian individuals in CaG and self-identified individuals of African ancestry in MHIB showed similar patterns (Figure 1B–C). In both cohorts, individuals were predominantly predicted as belonging to admixed African-Caribbean (ACB) or West African (YRI) populations. This concordance of predictions across biobanks suggests that a substantial portion of the selected MHIB individuals may have Haitian ancestry, which was confirmed with grand-parental information available in the metadata (Supplementary Results **S2.2**) and demonstrates that the approach successfully capture an admixed ancestry not explicitly present in the training set.

##### French Canadian population

The founder population of French Canadians (FC), the Western European majority group in both CaG and MHIB but absent from 1KGP, provided an opportunity to assess how the model generalizes to a well-represented but unmodeled population. FC individuals were typically predicted as CEUGBR or IBS (Figure 1B–C), illustrating the model’s capacity to interpolate across unrepresented but related groups.

Overall, across all examined populations, model predictions largely aligned with their continental ancestry, underscoring the generalizability of the model, while also capturing finer-scale patterns and uncertainties reflective of real-world population structure and labelling.

#### 4.3.2 Model predictions remain consistent on incomplete SNP data

We next examined the shift in model predictions when introducing missing genotype data at test time, a factor that can affect model generalizability such as when attempting to use the trained model on a new dataset which might not contain all the SNPs that were available during model training. We incrementally introduced missing data and compared model predictions to those obtained using complete SNP data. Figure 1D shows, for each investigated population in CaG and MHIB, the number of models whose predictions shifted relative to their outputs on complete data.

Prediction shifts varied by population: predictions for Italian individuals were the most consistent across the different levels of SNP missingness, while greater instability was observed among predictions for CaG Haitians, MHIB Africans, and French Canadians in both cohorts. This suggests that model predictions are more susceptible to missing values for admixed populations and populations lacking a closely related reference in 1KGP. Despite differences across populations, model predictions were generally robust to missing data: at 50% SNP missingness, at least 80% of predictions were consistent with those obtained using the complete SNP data. Moreover, the predicted populations generally remained within the same continental population with notable shifts occurring only near 99% missingness (Supplementary Figure 5). Examining prediction variability across increasing levels of missing data highlights the improved generalizability achieved through input dropout training, as models incorporating input dropout produced more consistent predictions across increasing amounts of missing genotypes than those trained without input dropout (Supplementary Figure 6).

#### 4.3.3 Hidden representations provide insights into unseen populations

The 1KGP dataset does not include any North African reference populations, while both CaG and MHIB include participants who self-identify as North Africans, including a subset of individuals in CaG reporting Moroccan ancestry. This presents a unique challenge: individuals from these populations have no close genetic counterparts among the training labels, limiting the model’s ability to assign meaningful predictions (Figure 2A). To better understand how the model handles such out-of-distribution samples, we examined the learned representations of these participants in the final hidden layer of the Diet Network models.

**Figure 2:**
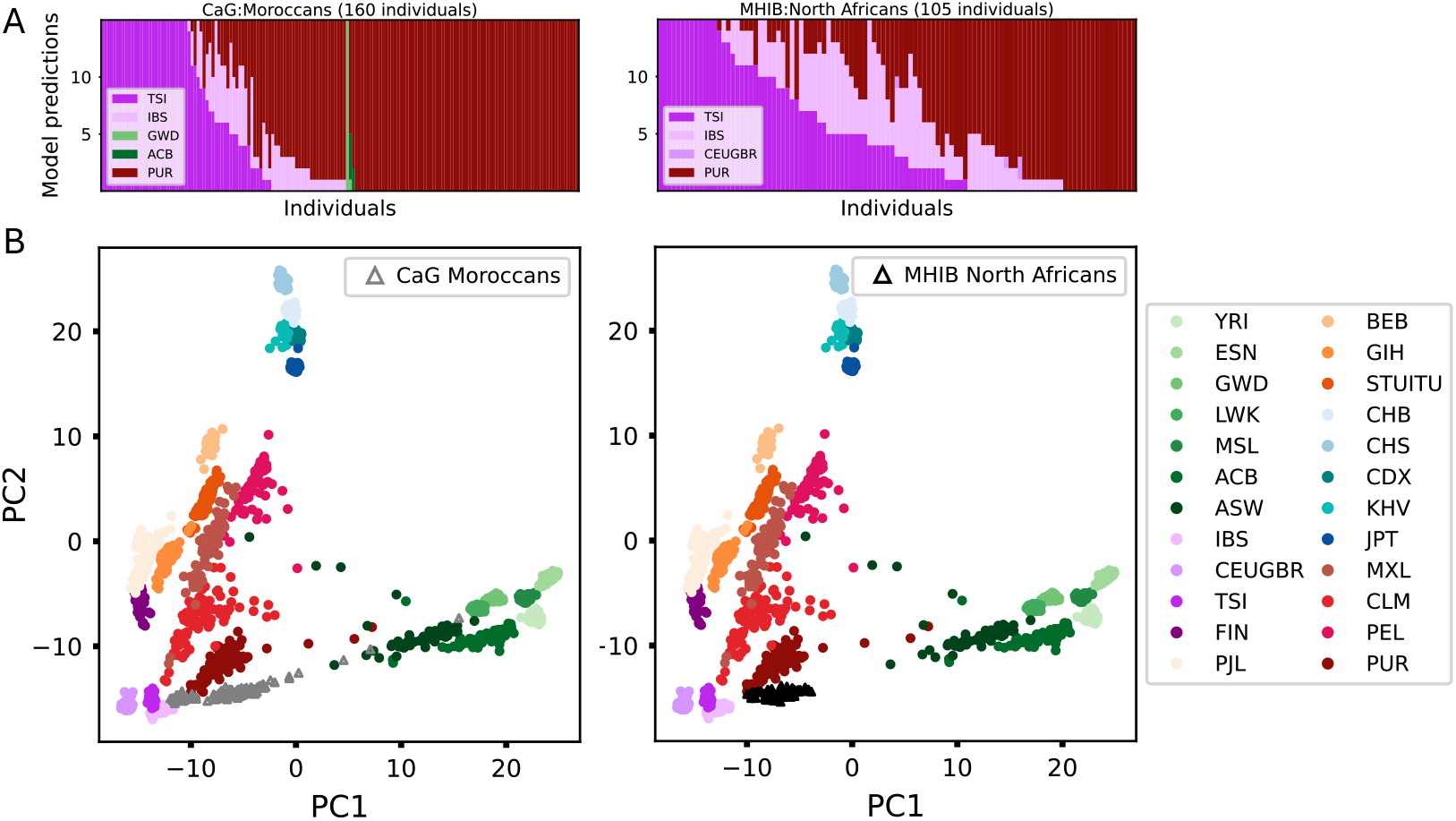
Diet Network hidden representation. **(A)** Predicted populations for individuals of Moroccan ancestry in CaG and North African ancestry in MHIB. Each vertical line represents an individual and the y-axis shows the population predicted by each of the 15 models. **(B)** Final layer PCA visualization for the 15 models based on 1KGP dataset with projected embeddings of Moroccan individuals in CaG and of North African individuals in MHIB.

We extracted the final hidden layer representations for 1KGP samples and performed PCA on these representations to construct a reference representation space. The same layer was extracted for CaG and MHIB North African individuals, which were then projected onto this PCA space, to visualize how the model positioned them relative to the 1KGP populations. These samples do not overlap with any specific 1KGP cluster but instead were positioned between individuals of EUR and AFR ancestries which aligns with their expected genetic ancestries (Figure 2B).

### 4.4 Uncovering Model-Leveraged SNPs Using Interpretability Methods

Understanding which genetic variants the Diet Network models rely on can help us evaluate how it makes predictions. We applied feature attribution gradient-based methods Saliency Maps [42], DeepLift [43], GradientSHAP [44] and Integrated Gradients [45], to estimate the importance of individual SNPs on the model’s predictions.

#### 4.4.1 Evaluating feature attribution methods

DeepLift, GradientSHAP and Integrated Gradients require a baseline input, which serves as a reference point to measure the effect of each SNP. Here, we use a genotype-average baseline, where each SNP is assigned the average genotype value across the training set. Because all inputs are standardized using z-score normalization before being passed to the network, this baseline input effectively becomes a vector of zeros, which can be interpreted as representing the absence of genetic information at each position.

The different gradient-based methods all produce local importance scores that we call local attribution scores. We evaluated whether these local attribution scores accurately capture the information used by the model by analyzing the impact of excluding high- and low-attribution SNPs on classification accuracy. For each individual, we ranked the SNPs using their local attribution scores computed with respect to their predicted population (see Methods). We progressively corrupted increasing numbers of SNPs based on their ranking, either starting with the highest scoring SNPs (ordered corruption) or the lowest scoring SNPs (reverse corruption). SNP corruption was performed by shuffling genotypes across individuals, referred hereafter as “shuffle corruption”. This approach disrupts the informative signal available to the model while preserving the overall genotype distribution.

For all attribution methods, the ordered corruption led to a faster decline in classification accuracy than corrupting the same number of randomly selected SNPs (Figure 3A), indicating that local attribution methods identified SNPs that are important to the model’s prediction. Conversely, reverse corruption resulted in a slower decline in accuracy, confirming that SNPs with low local attribution scores have little to no impact on the model’s predictions. Compared to the other attribution methods, the observed decline in accuracy for Saliency Maps was slower for the ordered corruption and faster for the reverse corruption, indicating that SNPs were ranked in a less optimal order.

**Figure 3:**
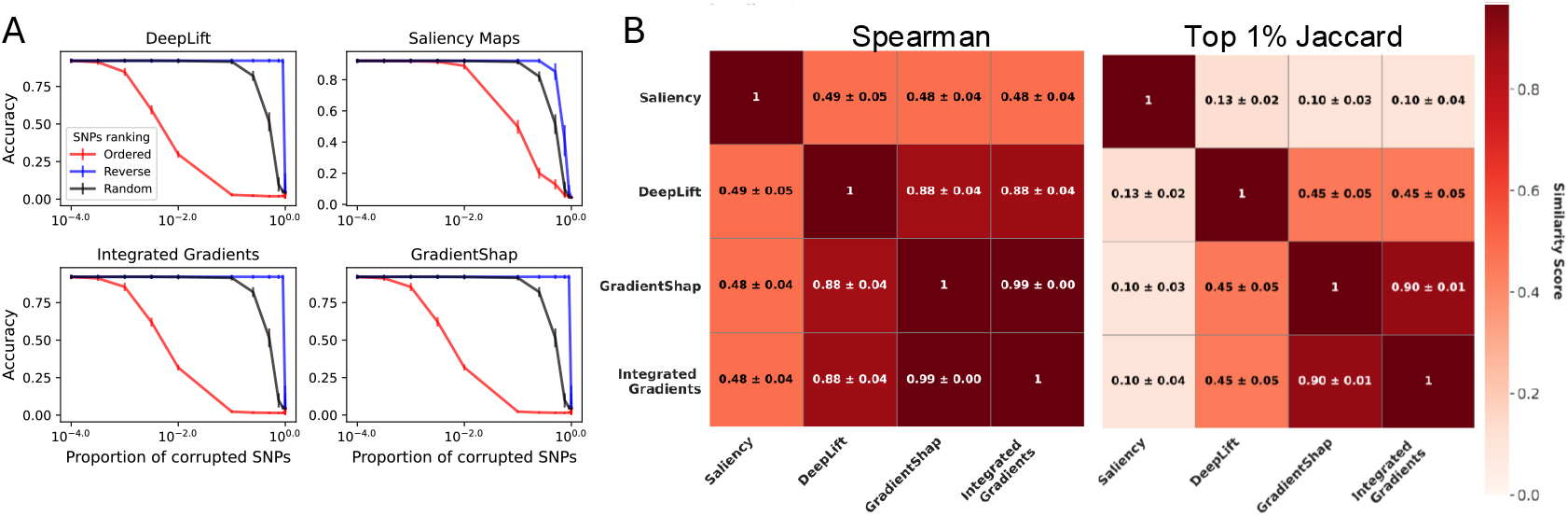
Performance of Feature Attribution Methods and Baselines. (A) Diet Network models classification accuracy on test sets for increasing proportions of corrupted features by feature attribution method. (B) Spearman correlation and top 1% Jaccard index comparing the similarity of attributions computed using between feature attribution methods (using the average-genotype baseline when applicable).

To quantify the decline in accuracy for each attribution method, we computed the area under the accuracy curves shown in Figure 3A. This resulted in two area metrics per method: the area under the ordered corruption curve (*A*_*ord*_) and the area under the reverse corruption curve (*A*_*rev*_).

The *A*_*ord*_ reflects the sensitivity of the attribution method, indicating to which degree the model relies on the top-ranked SNPs for its predictions. In contrast, the *A*_*rev*_ indicates the specificity of the attribution methods, assessing whether low-attribution SNPs truly have little effect on the model’s predictions. An ideal attribution method should yield a small *A*_*ord*_ and a large *A*_*rev*_. At a minimum, a reasonably effective attribution method should achieve a smaller *A*_*ord*_ and a larger *A*_*rev*_ compared to the area under the accuracy curve obtained when SNPs are corrupted in a random order (*A*_*ran*_). The area values (Table 1) confirmed that Saliency Maps consistently underperformed compared to the other methods in identifying SNPs that influence model predictions.

**Table 1:** Area metrics comparison of feature attribution methods

| Method | $A_{ord}$ | $A_{rev}$ |
| --- | --- | --- |
| DeepLift | $0.039 \pm 0.008$ | $0.877 \pm 0.013$ |
| GradientShap | $0.036 \pm 0.006$ | $0.878 \pm 0.013$ |
| Integrated Gradients | $0.036 \pm 0.007$ | $0.878 \pm 0.013$ |
| Saliency Maps | $0.201 \pm 0.020$ | $0.653 \pm 0.035$ |

*Area under the accuracy curves obtained with ordered SNPs corruption ( $A_{ord}$ ), reverse SNPs corruption ( $A_{rev}$ ) and random SNPs corruption ( $A_{ran} = 0.481 \pm 0.025$ ).*

To assess the consistency between attribution methods, we computed the Spearman correlation on the local attribution scores. Additionally, we used the Jaccard index on the top 1% of high-attribution SNPs to evaluate the overlap of important SNPs identified by different methods. The three top-performing methods, DeepLift, GradientSHAP, and Integrated Gradients, largely agreed on the SNP rankings (Figure 3B). As no single method consistently outperformed the others, we selected DeepLift for subsequent analyses as a representative, well-performing approach.

#### 4.4.2 Selecting an appropriate input baseline

We investigated how the choice of the input baseline influences the performance of the feature attribution method. Prior studies suggest that an effective baseline should resemble other samples in the dataset, reflect an absence of information and produce neutral predictions [45]. With this in mind, we designed three single-sample baselines tailored to our genetic context (detailed in the Methods section): (1) a genotype-average baseline, described above; (2) a reference baseline, homozygous for alleles in the human reference genome; and (3) a highest-entropy sample baseline, corresponding to the sample that yields the most uniform predictions across populations.

In addition to these single-sample baselines, we explored averaging contributions across multiple baselines, which can help reduce bias from any single point of comparison [46]. These included a set of randomly selected real samples and three synthetic sets, where genotypes were generated according to different distributions : (1) a uniform frequency set, where genotypes are drawn with equal probability 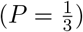;(2) a global frequency set, where genotypes are sampled based on their frequency in the dataset; and (3) a per-population frequency set, where genotypes are sampled per population according to that population’s genotype frequencies.

We evaluated the baseline performance using the *A*_*ord*_ and *A*_*rev*_ metrics under two corruption methods: the shuffle corruption method described above and a 1 Nearest Neighbor (1NN) imputation corruption method where genotypes are replaced by the genotypes of the nearest neighbor in the dataset (see Methods).

Among the single-sample baselines, the genotype-average baseline yielded the best performance, reflected by the lowest *A*_*ord*_ and highest *A*_*rev*_ values (Table 2). Among the multi-sample baseline sets, the randomly selected samples, global frequency, and per-population frequency baselines performed similarly, while the uniform frequency baseline performed worse. This likely stems from the fact that sampling genotypes uniformly, without adjusting for the actual genotype frequency distribution observed in the dataset, produces unrealistic inputs diverging from true individual genetic profiles. Furthermore, we observed that each single-sample baseline achieved the best performance when SNPs were corrupted by replacing their values with those of that same baseline, highlighting the importance of assessing baseline performance across multiple corruption methods (Supplementary Table 2). We also observed lower *A*_*ord*_ and *A*_*rev*_ performance when using supobtimal baselines, confirming that the area metrics can identify suboptimal baselines (Supplementary Figure 7).

**Table 2:** Area metrics comparison across baselines and corruption methods

| Baseline | Shuffle Corruption |  | 1NN Imputation |  |
| --- | --- | --- | --- | --- |
| | $A_{ord}$ | $A_{rev}$ | $A_{ord}$ | $A_{rev}$ |
| Reference | $0.049 \pm 0.008$ | $0.871 \pm 0.013$ | $0.046 \pm 0.006$ | $0.753 \pm 0.014$ |
| **Genotype-average** | $0.039 \pm 0.008$ | $0.877 \pm 0.013$ | $0.039 \pm 0.006$ | $0.84 \pm 0.014$ |
| Highest-entropy sample | $0.048 \pm 0.008$ | $0.875 \pm 0.013$ | $0.045 \pm 0.006$ | $0.796 \pm 0.016$ |
| Set - Random samples | $0.04 \pm 0.008$ | $0.877 \pm 0.013$ | $0.038 \pm 0.006$ | $0.839 \pm 0.014$ |
| Set - Uniform frequency | $0.055 \pm 0.008$ | $0.873 \pm 0.013$ | $0.051 \pm 0.006$ | $0.779 \pm 0.016$ |
| Set - Global frequency | $0.04 \pm 0.008$ | $0.877 \pm 0.013$ | $0.039 \pm 0.006$ | $0.839 \pm 0.014$ |
| Set - Per-population frequency | $0.04 \pm 0.008$ | $0.877 \pm 0.013$ | $0.038 \pm 0.006$ | $0.839 \pm 0.015$ |

Area under the accuracy curves obtained with ordered ( $A_{ord}$ ) and reverse ( $A_{rev}$ ) SNPs corruption. The genotype-average baseline was selected for downstream analyses.

The genotype-average baseline emerges as a reasonable and effective choice for input baseline. It consistently achieved the best or second-best performance across all corruption methods, while offering comparable results to the random, global frequency, and per-population frequency baseline sets, which are more computationally demanding.

### 4.5 Connecting Population Genetics Metrics with Feature Attributions

To further assess the biological relevance of the model’s attributions, we compared them to *F*_*ST*_, a well-established population genetics metric that quantifies allele frequency differentiation between populations. By using *F*_*ST*_ as a benchmark, we explored whether the SNPs identified as important by the model align with known patterns of population structure.

Population pairwise *F*_*ST*_ values, averaged across SNPs, are known to show a block structure with low genetic differentiation (*F*_*ST*_ value near 0) among populations from the same continent and higher differentiation between continents (Figure 4A). To assess whether attributions reflect population-differentiated SNPs, we computed a global importance score per SNP by averaging attribution values across individuals (see Methods) and used these scores to extract the highest-ranking SNPs in the dataset. Examination of *F*_*ST*_ values for the 10 SNPs with highest attribution SNP score showed strong differentiation patterns (*F*_*ST*_ values close to 1), with populations exhibiting pronounced divergence at these SNPs compared to 15 randomly selected SNPs, which displayed generally lower *F*_*ST*_ values (Figure 4B). These results suggest that the Diet Network model relies on SNPs that are differentiated between populations to make its predictions.

**Figure 4:**
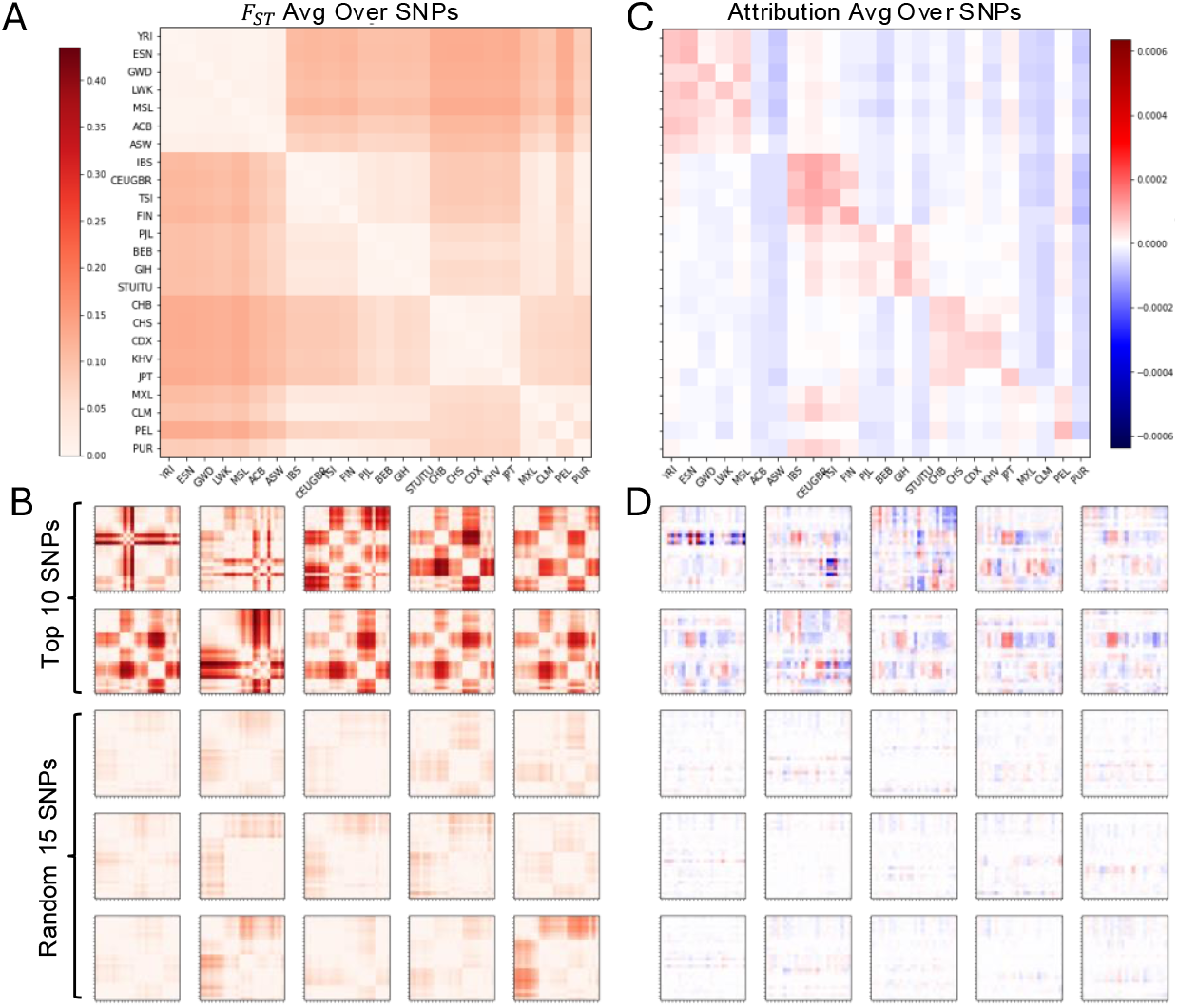
Comparing Attributions With Pairwise *F*_*ST*_ Scores. (A) 1KGP populations pairwise *F*_*ST*_ scores averaged across SNPs. (B) *F*_*ST*_ values of top 10 SNPs with the highest global attribution scores (top panel) and *F*_*ST*_ values of 15 randomly selected SNPs (bottom panel). (C) 1KGP populations pairwise attribution scores. Rows represent the true populations and columns indicate the candidate populations. The diagonal indicate the pairwise attribution score of correctly classifying individuals from a true population into the correct corresponding target population. (D) Pairwise attribution scores of the top 10 SNPs with the highest global attribution scores (top panel) and pairwise attribution scores of 15 randomly selected SNPs (bottom panel). These top and random SNPs are the same as in (B).

To explore how attribution scores reflect population structure, we computed a pairwise attribution matrix comparing actual and candidate population assignments (see Methods). The resulting heatmap shows a distinct block pattern along the diagonal, indicating that SNPs tend to contribute positively to predictions within the same continental group (Figure 4C). This structure closely mirrors that of the *F*_*ST*_ matrix (Figure 4A), suggesting that the model relies on population-differentiated signals to support population prediction. Within-population attribution scores are especially elevated for most populations, further confirming that the model accumulates evidence specific to each group for correct classification.

We found that model predictions for admixed populations behave differently: for four admixed populations, ACB, ASW, CLM and PUR, the attribution matrix showed low or negative values along the diagonal, suggesting that SNPs often reduced the probability of assigning individuals to their own population. These populations also consistently received negative attribution scores across all other groups, indicating a lack of strong positive evidence from the model. This pattern suggests that, for most populations, the Diet Network model assigns individuals by accumulating evidence based on SNPs characteristic of the population, whereas for the admixed ACB, ASW, CLM and PUR populations, assignment is driven by accumulating evidence against alternative populations.

When examining the pairwise attributions of the top 10 global attribution SNPs, their attribution patterns across populations mirror their *F*_*ST*_ profiles (Figure 4D). Specifically, SNPs that are not differentiated between an individual’s population and a similar target population tend to be used by the model as positive evidence for that target. Conversely, SNPs with high *F*_*ST*_ (high differentiation) relative to target populations are often used as negative evidence, reducing the probability of assignment to those targets. The similarity between the *F*_*ST*_ and attribution matrices is most evident for those SNPs with the highest attribution scores, suggesting that the model has learned to leverage population-differentiating signals in a way that reflects *F*_*ST*_ -captured population structure.

## 5 Discussion

The Diet Network is a deep learning model designed to handle genetic data and developed in the context of genetic ancestry prediction. Although the generalizability of the Diet Network trained on 1KGP has previously been evaluated [47] on another population reference dataset, the HGDP dataset, our study extends this work by assessing the model on two Canadian biobanks providing a practical assessment of generalizability. Furthermore, we gained insight into the model’s decision-making by applying feature attribution methods, which confirmed that the model relied on population-differentiated SNPs. This finding is supported by a comparison with pairwise *F*_*ST*_ values, a gold-standard metric in population genetics for quantifying genetic differentiation between populations. These interpretability experiments revealed that the Diet Network adopted two main prediction strategies: for most populations, it accumulated supporting evidence, while for specific admixed populations, it operated by eliminating alternatives, highlighting the complexity of admixed genomes. Additionally, we introduced a visualization of the model’s latent representations, enabling the characterization of individuals with admixed backgrounds not represented in the 1KGP reference panel used for training.

Several machine learning-based tools have been developed to infer genetic ancestry. For example, 23andMe employs a support vector machine (SVM) algorithm trained on a proprietary reference populational dataset [48, 49]. Their method performs local ancestry inference along the genome, producing ancestry proportions at varying levels of granularity depending on the algorithm’s confidence. However, this approach requires phased genotype data, which not only introduces potential errors or biases but also demands considerable domain expertise and computational resources, factors that may limit scalability and broad adoption beyond specialized research settings used by population geneticists. Another tool, SNVstory [30], uses two SVM models trained on 1KGP and the Simons Genome Diversity Project along with an XGBoost model trained on the gnomAD database. This ensemble method operate on whole-genome data to assign a single population label per individual at the continental level, with additional models used for finer-scale distinctions within continents. Yet, it remains unclear how such models handle admixed individuals or generalize to new datasets with differing SNP content.

A key challenge in applying these methods across datasets lies in the variability of available genetic markers, which inevitably differ between genetic studies, biobanks, and clinical cohorts due to diverse genotyping platforms or sequencing technologies and protocols. Inference methods that rely on a fixed reference panel or ancestry-informative markers (AIMs) [50, 51] are limited by their dependence on the presence of specific predefined SNPs, reducing their applicability for datasets lacking those markers. Furthermore, deep learning models trained on such fixed SNP sets may achieve high accuracy by over-relying on a narrow subset of highly informative variants, which can hinder generalizability.

To address these limitations, our method operates directly on unphased, unimputed genotype data, enhancing its adaptability to biobank and clinical datasets. Our training approach uses input dropout, encouraging the model to learn from a broader and more flexible set of SNPs by varying the subset it uses across samples, and mitigates overreliance on any fixed subset. As a result, the model remains robust to missing data, reducing the need for imputation and extensive preprocessing. We demonstrate that input dropout enhances the consistency of latent representations (Supplementary Figure 8), and reduces prediction variability across random weight initialization (Supplementary Figure 9). Moreover, unlike approaches requiring separate classifiers for different population panels, our method performs all predictions within a single unified model, simplifying deployment and improving scalability.

Despite their strong predictive performance and robustness to missing data, deep learning models like the Diet Network are often criticized as “black boxes”, offering limited transparency into the features driving their predictions. To address this, we applied feature attribution methods to uncover how the model leverages genetic information. Given the computational costs of permutation-based attribution methods, we employed gradient-based approaches, which are significantly more efficient, but require an appropriate input baseline as a point of comparison to quantify attributions. As most prior studies of feature attribution were developed in the context of computer vision, it remains unclear what constitutes an appropriate baseline for genetic data. We therefore designed and systematically evaluated multiple genomic baselines, and found that those most effective in our setting did not align with conventional criteria used in vision models, such as neutrality and uniformity of predictions (Supplementary Results **S2.3** and Supplementary Figures 10 and 11). This highlights the importance of developing domain-specific evaluation metrics, such as our proposed area-based measures, to rigorously assess baseline suitability across different genomic contexts and tasks.

Analyzing pairwise attribution scores between populations provided deeper insights into the classification strategies used by the model. The distinct attribution patterns observed for ACB, ASW, CLM, and PUR (Figure 4C) of negative or near-zero diagonal attribution scores, combined with consistently negative scores across all other population columns, suggests that the model struggles to identify shared, population-specific signals likely due to high inter-individual variability in ancestral contributions. In contrast, other admixed populations, such as PEL and MXL, showed stronger within-group attribution signals, suggesting the identification by the models of population-specific variants that enable more consistent classification.

The insights gained from these interpretability analyses also shed light on the behavior of the Diet Network in cases of uncertainty. Notably, the pairwise attribution score for correctly classifying PUR individuals (Figure 4C, where both the true and target population labels are PUR) was closest to zero in comparison to the other populations. This suggests that the model assigns individuals to the PUR population not because of strong supporting evidence, but rather due to the absence of strong evidence for alternative populations, effectively making PUR a “default” prediction in ambiguous cases. This behavior may explain why the PUR label was more frequently predicted when models were tested with high levels (99%) of missing data (Supplementary Figure 5). The PUR label was also assigned by a majority of models for the input baselines containing values near the genotype mean, effectively representing missing information due to input normalization (Supplementary Figure 11).

While effective in several respects described above, our approach has several limitations in its current form. First, it does not provide a built-in mechanism for estimating prediction uncertainty or confidence, a limitation in scenarios where misclassification risks must be explicitly quantified. At present, confidence could be approximated by examining the variability of model predictions for the same individual, both across multiple trained models and across the softmax probability distributions. While this approach does not yield formal confidence intervals or guarantees of correctness, as the model could consistently produce a consistent yet incorrect classification, it allows for the identification of individuals whose predictions are unstable or ambiguous. In addition, our visualization of latent representations enables the detection of individuals who are genetically distinct from the training populations, suggesting they may be out-of-distribution cases, thereby offering a complementary way to flag uncertain classifications.

At this stage, the Diet Network approach has not been implemented to support local ancestry inference, a feature offered by tools like RFMix [26] that assign ancestry at a finer genomic scale. Extending it to operate across genomic windows would enable local inference and could improve resolution in admixed genomes, a feasible direction for future work. Haplotype-level information could further enhance prediction accuracy by capturing linkage patterns that single SNPs may miss, although it would require rethinking the input representation or adapting the Diet Network to integrate hierarchical transformer architectures, which are well-suited for modelling haplotype-level information and long-range dependencies between genetic regions [52]. Although the Diet Network could be extended to predict complex phenotypes, the use of high input dropout, beneficial for generalizability, may hinder the model’s ability to learn meaningful patterns depending on the genetic architecture of the phenotype of interest. Therefore, model parameters such as dropout rates should be carefully tuned to the specific genetic context and goals of each application.

Accurate genetic ancestry prediction is essential for promoting equity in biomedical research and fairness in Artificial Intelligence (AI) tools for health. There is broad consensus on the need to develop predictive tools that are generalizable, unbiased and inclusive of global population diversity, but this effort depends on how ancestry is identified and annotated in health studies. Our method supports this goal by democratizing ancestry annotation within biobanks, including for individuals with missing data, by refining broad categories into more granular population groups.

Additionally, the method can facilitate the identification of individuals from local biobanks who are suitable for inclusion in population reference panels, thus leveraging the rich, region-specific genetic diversity available in these resources. As such, the Diet Network represents a step toward a more equitable foundation for AI development in health applications, supporting better assessment of model performance across ancestries in genomic research and ultimately supporting the design of fairer, more inclusive AI systems.

## 6 Methods

### 6.1 Data Processing

#### 6.1.1 1KGP dataset

The 1KGP whole-genome sequencing 30x coverage on GRCh38 dataset [37] was downloaded from the International Genome Sample Resource data portal. The dataset includes whole-genome sequencing at 30X coverage of 2,504 samples categorized into twenty-six populations across five continents (Supplementary Table 1). We merged closely related populations with minimal genetic differentiation: CEU and GBR (merged as CEUGBR), and ITU and STU (merged as STUITU), reducing the number of 1KGP populations from 26 to 24. These merges were based on low pairwise *F*_*ST*_ values and supported by population structure and relatedness analyses (further details and justification are provided in Supplementary Results **S2.1**). To mitigate class imbalance, CEUGBR and STUITU were downsampled randomly to match the largest population sample size within their respective continental groups (*n*_*CEUGBR*_ = 107; *n*_*STUITU*_ = 105).

#### 6.1.2 CaG dataset

We used data of 17,286 samples genotyped with the Illumina Infinium GSA v2 with Multi-Disease Panel and CaG_addon_v1_20037253_A2 genotyping array, as approved under projects 771235 and 406713. Population labels for FC individuals were assigned based on self-reported information: individuals had to identify as French Canadian, be of White European descent and have all four grandparents born in Canada. This resulted in 9,437 FC individuals. For Italian, Chinese, Haitian, and Moroccan ancestry groups, individuals were included if they reported all four grandparents born in Italy (367 samples), China (107 samples), Haiti (230 samples) and Morocco (160 samples).

### 6.1.3 MHIB dataset

We used data of 16,707 individuals genotyped on the GSA v3.0-MD genotyping array, as approved under project 2026-3546. Population labels were primarily assigned based on the information available in the self-reported ethnicity field as grand-parental information was less consistently available compared to CaG. These initial assignments were further refined through genetic analyses: principal component analysis (PCA) was used to identify a subgroup of 105 individuals with North African ancestry among those who had self-identified as Caucasian, and individuals whose genetic ancestry estimated using RFMix v2 [26] did not align with their self-reported ethnicity were excluded (Supplementary Results **S1.1**).

For the FC population, 12,884 individuals were included, who self-identified as Caucasian and reported that both parents were of French Canadian descent. The African ancestry group included 117 individuals who reported having ancestors originating from Africa or black ethnicity. The Asian ancestry group, included 62 individuals who self-identified as Asian. Additionally, we identified a subset of 419 individuals of Italian descent, based on consistent grand-parental information indicating that all four grandparents were born in Italy.

#### 6.1.4 Construction of an harmonized SNP set

The SNP set used to train the Diet Network models on 1KGP and test them on CaG and MHIB was obtained by first identifying the largest common set of SNPs shared between the CaG and MHIB genotyping arrays. To build this maximal SNP set, the following steps were independently applied to the CaG and MHIB datasets. Alleles were harmonized to the GRCh38 reference genome using Will Rayner’s strand alignment tool and the strand file GSAMD-24v3-0-EA 20034606 A1-b38 from https://www.strand.org.uk (last accessed on November 24th 2022). SNPs with allele frequency differences exceeding 20% compared to the TOPMed reference panel [53] were removed, as they likely result from incorrect genomic conversion. SNPs with a missing rate above 10% were excluded and monomorphic sites were removed. The final maximal SNP set consisted of 566,791 SNPs, representing the intersection of the filtered SNPs from both CaG and MHIB datasets.

The maximal SNP set shared between CaG and MHIB was further intersected with the 1KGP dataset and additional filtering was performed using PLINK v1.9. SNPs with more than 10% missing rate or a minor allele frequency below 5% in 1KGP were removed. Linkage disequilibrium pruning was applied using the parameters –indep-pairwise 50 5 0.5. SNPs located in the human leukocyte antigen region were excluded (chromosome 6, positions 28,510,121 to 33,480,577 on GRCh38). These filtering steps resulted in a final set of 229, 986 SNPs. Missing genotypes were subsequently imputed using ShapeIT5 [54] in cohort-based mode, without relying on an external reference panel.

### 6.2 Diet Network Architecture and Training

The Diet Network consists of two jointly trained neural networks: a main network that performs population classification from individual genotype data, and an auxiliary network designed to reduce overfitting that generates SNP-specific weights for a high-dimensional input layer using a SNPs embedding, a vector representation of every SNPs (Supplementary Figure 3). As the original implementation [28] was based on the now-deprecated Theano library, we reimplemented the architecture in PyTorch. The code is available at https://github.com/HussinLab/DIETNETWORK.

We trained 15 Diet Network models on the 1KGP dataset (2,322 individuals from 24 populations) using 5-fold cross-validation, repeating the process three times. Each fold was split into training (60%), validation (20%), and test (20%) sets, with population labels balanced between the splits. Genotype input features were standardized by subtracting the mean and dividing by the standard deviation calculated on genotypes from the training set. For the auxiliary network, SNP embeddings were defined as genotype frequencies per population in the training set. Hyperparameter tuning was conducted on the validation set. Further details on model training are provided in Supplementary Methods **S1.2**. The trained model weights, the accompanying files required to run the models and the 1KGP dataset used to train the models are available at https://doi.org/10.5281/zenodo.16943453.

### 6.3 Generalizability Analyses

To assess the generalizability of the Diet Network, we performed both in-sample (1KGP) and out-of-sample (CaG and MHIB) evaluations under varying levels of missing genotype data. In 1KGP, we simulated SNP missingness by removing predefined proportions of input features and compared classification accuracy across models trained with different input dropout rates (Supplementary Methods **S1.2**). Models trained with a 99.5% input dropout rate were applied to CaG and MHIB datasets, with prediction variability reported across fifteen models per individual. Prediction shifts were tracked across varying proportions of missing SNPs compared to using complete SNP data (Supplementary Methods **S1.2**). In all experiments, the genotypes of the remaining SNPs were rescaled to account for the reduced input dimensionality resulting from SNP removal (Supplementary Methods **S1.3**).

To examine how individuals are represented by the Diet Network models, we extracted activations from the last hidden layer (100 neurons) across all 15 trained models, resulting in a 1,500-dimensional representation per individual. These latent representations provide a compressed encoding of the input data learned by the network and can be extracted for any individual. To examine the learned population structure and potential out-of-distribution signals, we performed PCA (Scikit-learn 1.0.1 [55]) on the combined 1,500-dimensional latent vectors from all 1KGP individuals and projected the latent vectors of CaG Moroccan and MHIB North African individuals onto this PCA space. The resulting principal components enabled joint visualization of how these groups relate to the training populations in the learned representation space.

### 6.4 Interpretability Framework and Baselines

We applied four feature attribution methods using the Captum library: Saliency Maps, Integrated Gradients, DeepLIFT, and GradientSHAP. Each attribution method outputs local scores with dimensions SNP *×* population *×* individual, quantifying the contribution of each SNP to each predicted population class. For attribution methods requiring a baseline input (all except Saliency Maps), we designed several biologically informed single-sample and multi-sample baselines. All baselines matched the input dimensionality used to train the Diet Network models (229,986 SNPs). Single-sample baselines consisted of a single vector of this length, defined as follows:

- *Genotype-average*: Each SNP was assigned the mean genotype value computed across the training set. After input z-score normalization (Supplementary Methods **S1.2**), this baseline corresponds to a zero vector, effectively representing a sample with all genotypes missing.
- *Reference*: Each SNP was set to the homozygous reference allele as defined in the GRCh38 reference genome. For biallelic SNPs, this corresponds to a genotype of 0.
- *Highest entropy sample*: The individual from the training and validation sets whose model prediction yielded the highest entropy (i.e., most uniform probability distribution across the 24 1KGP population classes) was selected.

Each multi-sample baseline set was composed of 96 vectors:

- *Random samples set* : 96 individuals randomly selected from the training and validation datasets.
- *Uniform set* : 96 synthetic samples where each SNP position was assigned genotype values 0, 1, or 2 with equal probability 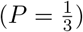.
- *Global set* : 96 synthetic samples where genotypes were drawn based on the global genotype frequencies observed across all 1KGP individuals.
- *Per-population set* : 96 synthetic samples where genotypes were drawn according to population-specific genotype frequencies from the 1KGP dataset, with 4 samples per population.

For all multi-sample baselines, attribution scores were computed individually for each sample in the set and then averaged [56].

For each attribution method and each baseline, this resulted in a total of 229, 986 *×* 24 local attribution scores per individual. These scores reflect, for each individual, the contribution of a given SNP to the predicted probability of every 1KGP population. Scores were computed for individuals in the test set of each of the 15 Diet Network models.

### 6.5 Global and Pairwise Attribution Scores

We aggregated local attribution scores to provide a single importance value per SNP across individuals. To compute these global attribution scores, we first selected the local attribution score computed with respect to the class predicted by the model for each individual. We then averaged these selected scores across individuals to obtain one global attribution score for each SNP. We evaluated alternative aggregation strategies (Supplementary Methods **S1.4**, Supplementary Figure 12 and Supplementary Table 3), but simple averaging provided the most robust results for identifying model-informative SNPs.

We also computed pairwise attribution scores between true and target populations, reflecting how SNPs contribute to misclassification across groups. To compute these pairwise scores, individuals were first grouped by their true population. For each target population, we averaged the local attribution scores across all SNPs, yielding a single value per true-target population pair.

### 6.6 Benchmarking Attributions Using SNP Corruption and Population Differentiation

To assess whether attribution methods effectively identify important SNPs, we progressively corrupted input genotypes of individuals in the test sets based on attribution scores and measured the resulting change in classification accuracy. For each individual, SNPs were ranked according to their local attribution scores relative to the model’s predicted class, and were corrupted in descending, ascending, or random order of importance using several corruption strategies.

We investigated several strategies for corrupting SNPs using DeepLift attribution scores.

- *Shuffle corruption*: For each corrupted SNP, genotype values were randomly permuted across the test set. This preserves the genotype distribution at each SNP position while disrupting individual-specific genomic patterns.
- *1NN imputation*: Corrupted SNPs were replaced with the corresponding genotypes from the individual’s nearest neighbor, identified using Euclidean distance on non-corrupted SNPs from the training and validation sets.
- *Genotype-average corruption*: Corrupted SNPs were replaced by the corresponding values in the genotype-average baseline.
- *Reference corruption*: Corrupted genotypes were replaced by those from the Reference baseline.
- *Highest-entropy sample corruption*: Corrupted values were drawn from the Highest entropy sample baseline.

For the genotype-average and reference corruption strategies, uncorrupted SNPs values were rescaled (Supplementary Methods **S1.3**).

To quantify the performance of attribution methods, the area under the accuracy curves was computed. These curves reflect how model accuracy changes as an increasing proportion of SNPs are corrupted. This evaluation yields two area metrics: *A*_*ord*_, corresponding to corruption in descending order of importance (from highest to lowest local attribution score), and *A*_*rev*_, for corruption in ascending order (from lowest to highest local attribution score).

Pairwise *F*_*ST*_ values were computed between all pairs of the 24 populations for all 229, 986 SNPs from the harmonized SNPset using VCFtools 0.1.16. Pairwise *F*_*ST*_ matrices were generated for top SNPs identified with global attribution scores and randomly selected SNPs to assess correspondence between genetic differentiation and model-derived importance.

## 7 Declarations

## 7.1 List of abbreviations

1KGP: Thousand Genomes Project
1NN: 1 Nearest Neighbor
A_ord_: Area under the accuracy curve obtained when corrupting SNPs in descending order of attribution score
A_rev_: Area under the accuracy curve obtained when corrupting SNPs in ascending order of attribution score
A_ran_: Area under the accuracy curve obtained when corrupting SNPs in random order
AI: Artificial Intelligence
AIM: Ancestry Informative Marker
CaG: CARTaGENE
CEUGBR: Population label of CEU and GBR individuals
FC: French Canadian
F_st_: Fixation index (a measure of population differentiation)
GSA: Global Screening Array
HGDP: Human Genome Diversity Project
MHIB: Montreal Heart Institute Biobank
PCA: Principal Component Analysis
SNP: Single Nucleotide Polymorphism
STUITU: Population label of STU and ITU individuals
SVM: Support Vector Machine

## 7.2 Ethics, consent and permissions

We obtained ethics approval for this study from the Montreal Heart Institute Institutional Review Board (IRB).

## 7.4 AI tools disclosure

During the preparation of this work the authors used Large Language Models to improve the readability and language of sections of the manuscript. After using these tools, the authors reviewed and edited the content as needed and take full responsibility for the content of the publication.

## 7.6 Author’s Contributions

**C.R.-B** and **M.S** : Conceptualization, Formal analysis, Investigation, Methodology, Software, Validation, Visualization, Writing – original draft, review & editing; **R.P** and **J.-C.G**: Data curation, Formal analysis, Methodology, Validation, Visualization, Writing – review & editing; **PL.C**: Validation, Writing – review & editing; **S.L**: Supervision, Validation, Writing – review & editing; **J.GH**: Conceptualization, Funding acquisition, Project administration, Resources, Supervision, Validation, Writing – review & editing

## 8 Acknowledgements

We thank Yoshua Bengio, Marc-André Legault, and Olivier Tastet for their valuable discussions and insights regarding the experiments and results. This work was made possible through computational resources provided by the Digital Research Alliance of Canada.

