## Supplementary Material for "A Transparent and Generalizable Deep Learning Framework for Genomic Ancestry Prediction"

#### S1 Supplementary Methods

##### S1.1 Self-Reported Ethnicity Labels in Biobank Cohorts

Evaluating model generalizability across datasets in population genetics is complicated by context-dependent population labeling of individuals in biobanks. While the Thousand Genomes Project (1KGP) populational dataset uses well-defined inclusion criteria, biobank data typically rely on self-reported ethnicity, which can vary in accuracy and resolution as it relates to genetic ancestry. This mismatch introduces uncertainty in defining ground truth labels, complicating the assessment of model generalizability. To navigate this, we implemented strategies to assign population label tailored to the available metadata in CaG and MHIB. It is important to note that the terminology and classification schemes used in these questionnaires were defined by the original biobank protocols and reflect prevailing standards at the time of data collection.

###### S1.1.1 CaG

Participants in the CaG cohort completed a self-administered socio-demographic and lifestyle questionnaire, which included questions on ethnicity and familial origin. Ethnicity was self-reported by selecting one or more of the following predefined categories: White European, Black, Arab, Latino, Southeast Asian, East Asian, West Asian, South Asian, Jewish, or Other. In addition, individuals were asked to provide information on their country of birth, as well as the countries of birth of their parents and grandparents.

Population labels were derived from the reported birth countries of participants' grandparents. These populations included individuals of Italian ancestry, closely aligned with the *Toscani in Italy* (TSI) population from 1KGP, and individuals of Chinese ancestry, who may correspond to several 1KGP reference groups, including *Han Chinese in Beijing* (CHB), *Han Chinese South* (CHS), and *Chinese Dai in Xishuangbanna* (CDX). We also evaluated model performance on individuals with African-European admixed ancestry, focusing on Haitian individuals, a population with known admixture but not represented in 1KGP. Nevertheless, similar admixed populations such as *African Caribbeans in Barbados* (ACB) and *African*

*Ancestry in the Southwest US* (ASW) are included in 1KGP. Furthermore, we assessed predictions for French Canadians (FC), a population of European descent that constitutes the majority of the biobank but is absent from the 1KGP reference. This offered a valuable test case for evaluating model generalization to a well-sampled yet unrepresented population. Lastly, we included a population entirely absent from 1KGP, the Moroccan population, representing North-African ancestry.

##### S1.1.2 MHIB

Participants in the MHIB cohort self-reported their ancestry by selecting from predefined categories that included: Caucasian (defined as having ancestors from Europe, North Africa, or the Middle East), Hispanic (e.g., Mexican, Puerto Rican, Cuban, South or Central American), Black (ancestors from African or Black ethnicity), Asian (ancestors from the Orient, Asia, Indian subcontinent, or Pacific Islands—including China, Japan, Philippines, Korea, and Samoa), Native American (ancestors from any North American Indigenous group or tribe), and Other. These labels and definitions reflect the original wording used in the MHIB questionnaire and associated metadata. We recognize that some of these terms may be considered outdated or sensitive due to their cultural and historical connotations. Their inclusion here is intended solely to preserve the integrity of the source data, and does not imply endorsement of these classifications.

As in CaG, individuals of Italian ancestry in MHIB were identified based on grandparental birth country information. However, due to less consistent documentation of grandparental information in MHIB, we relied on the self-reported ethnicity field to define labels for other population groups. For our analysis, we retained individuals of Asian ancestry, a broad category that can relate to both East Asian and South Asian populations in 1KGP, African ancestry, and French Canadian descent.

To ensured that our evaluation focused on samples with reasonably consistent ancestry information, we further improved the accuracy of population assignments derived from the self-reported ethnicity field by applying additional filtering steps based on genetic ancestry. North African ancestry individuals were identified using principal component analysis (PCA) with FlashPCA2 [1], which revealed a genetically distinct cluster within individuals who had self-identified as Caucasian. This subgroup was reclassified as North African, in accordance with the questionnaire structure that includes North African origins within the Caucasian category. We further filtered the dataset using genetic ancestry inference using RFMix v2 [2], removing individuals whose inferred global genetic ancestry did not align their self-reported category. RFMiX analysis was conducted using individuals from the 1KGP representing European (EUR), African (AFR), East Asian (EAS), and South Asian (SAS) superpopulations (excluding the admixed ACB and ASW populations) as the reference panel. Based on global ancestry proportions, the following individuals were excluded:

- 1 individual who self-identified as French Canadian (FC) but had an EUR ancestry proportion $< 0.7$;
- 9 individuals who self-identified as African but had an AFR ancestry proportion $< 0.7$;
- 4 individuals who self-identified as Asian but had an EAS or SAS ancestry proportion $< 0.7$.

#### S1.2 Details of Diet Network training and cross-dataset validation

##### S1.2.1 Training on 1KGP data

Fifteen Diet Network models were trained on the 1KGP dataset using 5-fold cross-validation, each fold repeated across three different random seeds to assess variability due to random initialization. The input dataset consisted of 2,322 individuals across 24 merged population labels. Each fold was split into training (60%), validation (20%), and test (20%) sets, ensuring that every individual appeared once in the test sets across folds. Population labels were balanced between training, validation and test splits.

For the auxiliary network, SNP embeddings were derived from genotype frequencies computed from the training set within each fold (see Supplementary Figure 3). Genotype inputs to the main network were standardized by subtracting the mean and dividing by the standard deviation of each SNP, calculated across the training set.

The number of layers, neurons per layer, optimizer and loss function were kept consistent with the original Diet Network implementation [3]. A hyperparameter search was conducted to optimize the learning rates for the main and auxiliary networks using performance on the validation set.

We trained multiple models using a range of input dropout rates (0, 0.5, 0.9, 0.99, 0.9925, 0.995, 0.997, 0.999, 0.9995, 0.9999), and identified the optimal input dropout rate by evaluating classification accuracy on test sets with varying proportions of randomly masked SNPs (0.1, 0.25, 0.5, 0.7, 0.8, 0.9, 0.95, and 0.99). SNP masking was performed as follows: for missingness proportions up to 0.5, we generated non-overlapping SNP subsets to ensure each SNP was removed at least once (e.g., two non-overlapping subsets for a missingness proportion of 0.5, 100 non-overlapping subsets for a proportion of 0.01). To ensure consistent comparisons, we generated 100 subsets for each missingness proportion. For missingness proportion above 50%, subsets were instead constructed to ensure that each SNP was included at least once. Genotypes of the remaining SNPs were rescaled to account for reduced input dimensionality using a scaling factor based on the ratio of SNPs in the training and test sets (Supplementary Methods **S1.3**).

##### S1.2.2 Testing on 1KGP data

The 1KGP dataset was split to ensure each individual appeared in the test set exactly once across the 5 cross-validation folds. The overall performance was evaluated using predictions from the five models specific to each fold. To assess the impact of model stochasticity, the 5-folds cross-validation experiment was repeated across three different random seeds, resulting in a total of 15 models. Consequently, each individual was evaluated by three models, enabling analysis of variability due to random initialization. Model predictions are presented in Supplementary Figure 4.

##### S1.2.3 Cross-dataset validation on CaG and MHIB

To evaluate generalizability, each individual from the CaG and MHIB datasets was tested using the 15 Diet Network models trained on 1KGP data with 99.5% input dropout. This produced fifteen population label predictions per individual. Although all SNPs used for model training were present in the CaG and MHIB datasets, some genotypes were missing at the individual level (no more than 10% per SNP). These missing genotypes were imputed with the mean SNP value from the corresponding 1KGP training set for each fold, which corresponds to a value of 0 after input feature standardization.

To further assess the impact of missing SNPs on model generalizability, we performed simulations by randomly removing SNPs at different proportions of missingness (0.25, 0.50, 0.75, 0.90, 0.95, and 0.99) from the CaG and MHIB datasets. For each proportion, one random subset of SNPs was removed, and the resulting predictions were compared to those obtained on the complete SNP data. Genotypes of the remaining SNPs were rescaled to account for reduced input dimensionality using a scaling factor based on the ratio of SNPs in the training and test sets (Supplementary Methods **S1.3**).

#### S1.3 Scaling Missing and Corrupted Inputs

##### S1.3.1 Preliminaries and Notation

Let $X = (x_1, ..., x_D)$ be a vector of size $D$, where each element $x_i \in (0, 1, 2)$ represents the genotype at a single SNP $i$. Prior to being fed into the network, each genotype $x_i$ in $X$ is standardized using the mean $\mu_i$ and standard deviation $\sigma_i$ of the corresponding SNP, calculated from the training data. Let $Y = (y_1, ..., y_D)$ denote the vector of standardized genotypes where $y_i = \frac{x_i - \mu_i}{\sigma_i}$.

Let $C$ denote the set of corrupted positions, let $Y' = (y'_1, ..., y'_D)$, denote the vector of data where $y'_i = y_i$ when $i \notin C$ and $y'_i$ is corrupted when $i \in C$. We want to scale up $Y'$ in order to keep $\sum y'_i$ as close as possible as $\sum y_i$ Assuming that:

$$\sum y_i \approx \frac{D}{|C|} \sum_{i \in C} y_i \approx \frac{D}{D - |C|} \sum_{i \notin C} y_i \tag{1}$$

##### S1.3.2 Scaling for missing inputs and Genotype-average corruption method

We define missing data or data corrupted with the Genotype-average corruption method $Y'$ such that if position $i$ is missing or corrupted, then $y_i$ is replaced by 0. We define $Y'$ as:

$$y'_i = \frac{D}{D - |C|} \left\{ \begin{array}{ll} y_i, & i \notin C \\ 0, & i \in C \end{array} \right\} \tag{2}$$

By scaling our data by $\frac{D}{D-|C|}$ we can then make $\sum y'_i$ closer to $\sum y_i$.

$$\sum y'_i = \frac{D}{D - |C|} \sum_{i \notin C} y_i \approx \sum y_i \tag{3}$$

##### S1.3.3 Scaling for reference corruption method

If our corruption involves replacing the value with the reference, then corrupting position $i$ results in $y_i$ being replaced by $-\frac{\mu_i}{\sigma_i}$.

A similar computation yields the scaling and translating needed. If we let $\mu$ and $\sigma$ be the average mean and standard deviation of the $\mu_i$ and $\sigma_i$ respectively, we define $Y'$ as:

$$y'_i = \frac{D}{D - |C|} \left\{ \begin{array}{ll} y_i & i \notin C \\ -\mu_i/\sigma_i & i \in C \end{array} \right\} + \frac{D}{|C|}\frac{\mu}{\sigma} \tag{4}$$

By scaling our data by $\frac{D}{D-|C|}$ and translating it to $\frac{D}{|C|}\frac{\mu}{\sigma}$ we can then make $\sum y'_i$ closer to $\sum y_i$.

$$\sum y'_i = \frac{D}{D - |C|} \sum_{i \notin C} y_i + \frac{D}{|C|}\frac{\mu}{\sigma} \approx \sum y_i \tag{5}$$

#### S1.4 Aggregating Local Attribution Scores into Global Attribution Scores

We use four feature attribution methods in this study, that provide local attribution scores per SNP and per individual: Saliency Maps, Integrated Gradients, DeepLIFT, and GradientSHAP. Briefly, Saliency Maps compute the gradient of the model output with respect to the input features [4]. Integrated Gradients estimate feature importance by integrating the gradients of interpolated values along a linear path between the input and one or more baselines. DeepLIFT calculates the difference in neuron activations between the input and a reference baseline, backpropagating these contribution scores through the network. GradientSHAP estimate feature importance by averaging gradients along randomly sampled paths between the input and a baseline, estimating Shapley value based on these integrated gradients.

We consolidated the local attribution scores returned by the attribution methods into a single global attribution score per SNP using several aggregation strategies, as there is currently no established consensus in the machine learning literature for converting local attributions to global importance. Specifically, we compared using average and rank average of local attribution values, with the latter aiming to reduce the influence of outliers. Additionally, we explored averaging local attribution scores across individuals grouped by either their true population or predicted population, in order to capture population-specific signal, acknowledging that certain SNPs may only be highly attributed within specific populations.

We performed a series of SNP corruption experiments in which we corrupted increasing number of SNPs based on their ranking derived from global attribution scores. In this setting, the same set of SNPs was removed for all individuals, in contrast to corruption experiments based on local attribution rankings, where the set of SNPs removed varied between individuals.

The area $A_{ord}$ and $A_{rev}$ metrics revealed that averaging local attributions yielded better performance than the other aggregation strategies tested (Supplementary Figure 12 and Supplementary Table 3). Specifically, SNP corruption based on global attribution rankings yielded lower $A_{rev}$ values compared to random SNP corruption under the "replace with missing" corruption method. This result suggest that, in this case, less informative SNPs were more effectively identified through random selection than by feature attribution. However, this pattern was not observed with the resample value and 1NN substitution corruption methods. We concluded that the observed worst performance of corrupting low attribution

SNPs compared to random under the "replace with missing" corruption is more plausibly attributed to the stronger impact of missing values on model predictions.

### S2 Supplementary Results

#### S2.1 Population Label Merging in 1KGP Dataset

We merged the CEU (Utah residents with Northern and Western European ancestry) and GBR (British from England and Scotland) populations under a single population label (CEUGBR), as they share recent ancestry from the same region of Europe, despite being geographically distinct, as done previously [5]. Among all European population pairs in 1KGP, CEU-GBR exhibited the lowest pairwise $F_{ST}$ value (0.0003), computed with VCFtools 0.1.16 across the $229,986$ SNPs from the shared SNPset, compared to other European pairs (0.0015 to 0.0119) and across the entire dataset (0.0008 to 0.0178).

Among South Asian populations, within-population substructure was observed, with subgroups showing greater genetic similarity to samples from other SAS populations than to their own population [6]. Specifically, ITU (Indian Telugu in the UK) and STU (Sri Lankan Tamil in the UK) were merged under a single population label (STUITU) as they had the lowest $F_{ST}$ values (0.0012) compared to the other pairwise $F_{ST}$ of SAS populations (0.0022 to 0.0044). PCA of these two populations (Supplementary Figure 2) revealed two distinct clusters within the ITU group, with one cluster genetically closer to the STU population than to other ITU individuals. Furthermore, a relatedness analysis identified a pair of siblings split between ITU and STU populations [7], further supporting the merging of these two populations.

#### S2.2 Grand-parental information of MHIB Africans

Of the 117 individuals with self-reported African ancestry in MHIB, grand-parental country of birth data were available for 58, with 46 reporting at least one grandparent born in Haiti. This suggests that a substantial proportion of self-reported African individuals in MHIB may be from Haiti, which could explain the consistency in model predictions observed between the CaG and MHIB cohorts. Furthermore, compared to the prediction for Haitians in CaG, there was broader representation of other African populations (e.g., LWK, GWD, ESN, MSL) in MHIB.

#### S2.3 Investigating Baselines Neutrality

We evaluated whether our genomics-informed baseline inputs satisfy prediction neutrality, the expectation that model predictions for baseline inputs should not be biased toward specific classes (populations). This assessment was conducted for both single-sample baselines and sets of baselines using two metrics: (1) the entropy of the predicted probability distributions returned by the models, which reflects the uncertainty of the predictions, and (2) the variability of predicted classes across models, which indicates how evenly predictions are distributed across categories. We assessed the consistency between the best-performing baselines identified using the $A_{ord}$ and $A_{rev}$ metrics (i.e., the genotype-average single baseline and the multi-sample baselines: random sets, global set, and per-population set) and those characterized as more neutral based on our entropy and class diversity criteria.

###### S2.3.1 Entropy

Among all baselines, the Highest-entropy single-sample baseline and the Uniform frequency baseline set exhibited the highest prediction entropy (Supplementary Figure 10), despite being among the worst performers based on the area metrics (Main Table 2). Conversely, the random samples and per-population set baselines, which achieved high performance according to the area metrics, displayed the lowest prediction entropy. This suggests that prediction neutrality, as measured by entropy, does not reliably distinguish between effective and ineffective baselines in our genomic context.

###### S2.3.2 Prediction variability

Investigating the prediction variability revealed that the per-population baseline set, one of the top performers based on the area metrics (Main Table 2), was the only baseline to produce a balanced distribution of predicted classes across all populations (Supplementary Figure 11). In contrast, other high-performing baselines, such as the Genotype-average and Global frequency set exhibited prediction biases toward specific populations, particularly PUR. This indicates that prediction neutrality, as measured by the variability in the predicted classes, does not consistently identify effective baselines.

Overall, the assessment of baseline neutrality using entropy and predicted class variability does not provide reliable guidance for determining the suitability of baselines in our genomic context, emphasizing the importance of incorporating our area metrics in the evaluation process.

### S3 Supplementary Figures

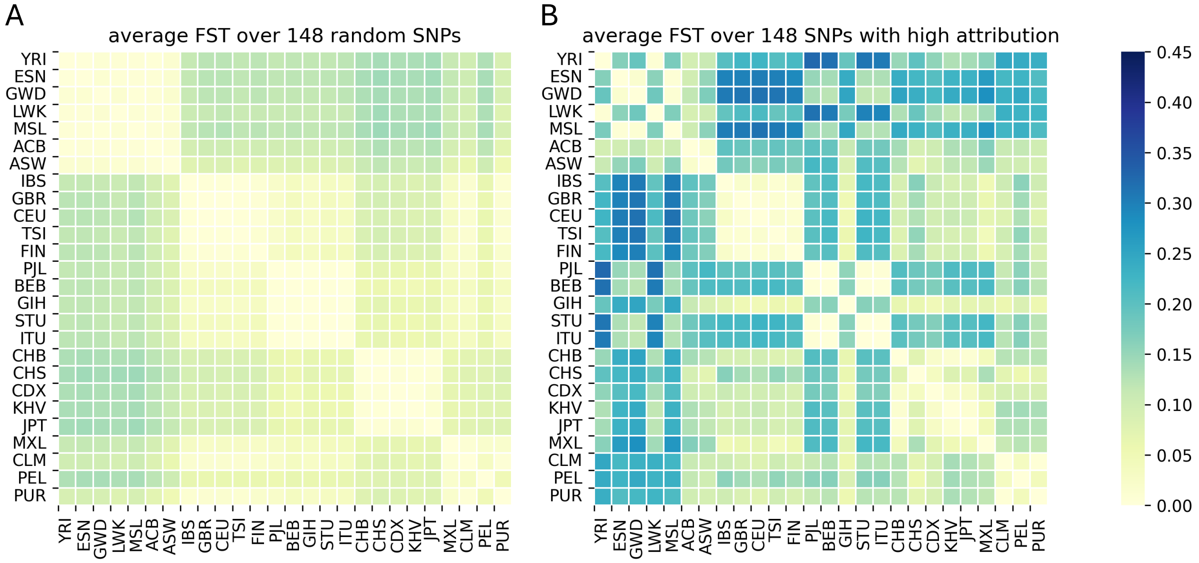


**Supplementary Figure 1: Population pairwise $F_{ST}$ of random and highly attributed SNPs.** Pairwise $F_{ST}$ values computed between all 1KGP population pairs and averaged across SNPs using data from the 1KGP Phase 3 dataset which includes 3,450 individuals genotyped in batches using the Affymetrix genotyping array. SNPs were filtered following the steps outlined in the Methods section, yielding 294,427 SNPs. (A) Average pairwise $F_{ST}$ of 148 randomly selected SNPs shows typical population structure patterns, with populations within continental groups having lower $F_{ST}$ values. (B) Average pairwise $F_{ST}$ for the 148 SNPs with the highest attribution scores identified using the Integrated Gradients method applied to fifteen Diet Network models trained via 5-fold cross-validation with three random seeds. The $F_{ST}$ patterns of those high attribution SNPs exhibit unexpected divergence. Notably, unusually high differentiation was observed between pairs of closely related populations including YRI with ESN, GWD, and MSL, and GIH with PJL, BEB, ITU, and STU. These SNPs showed unexpected divergence, including high differentiation between closely related populations (e.g., YRI vs. ESN/GWD/MSL; GIH vs. PJL/BEB/ITU/STU). Comparing 1KGP genotyping array data to the 1KGP high-coverage whole genome sequencing data revealed allele frequency discrepancies greater than 20%, suggesting genotyping errors not removed by standard filters.

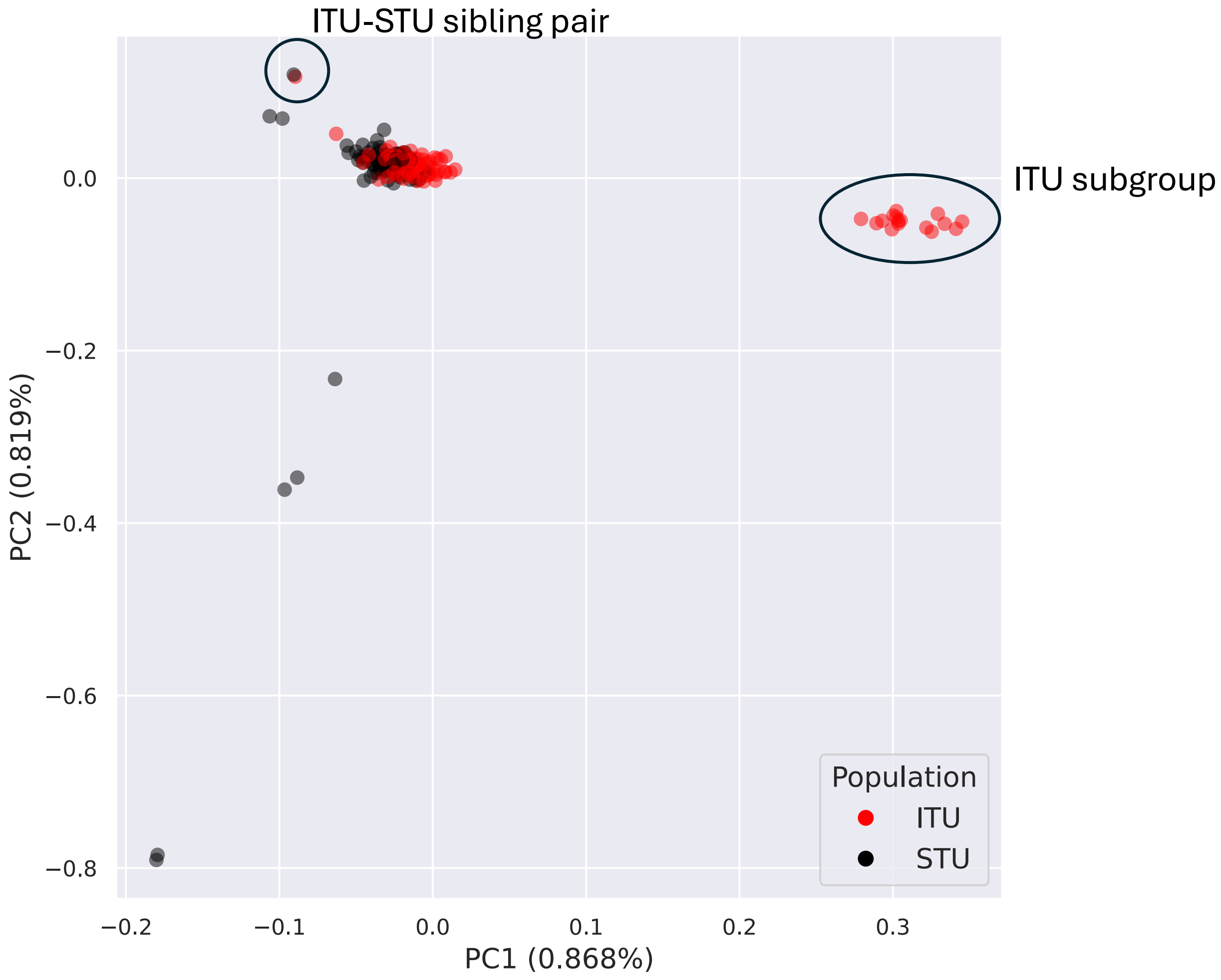


**Supplementary Figure 2: PCA of ITU and STU populations in 1KGP shows ITU subgroup and STU-ITU sibling pair.** PCA was performed using 204 samples from ITU and STU populations and 230K SNPs used to train the Diet Networks models. PCA shows two distinct clusters within the ITU group, with one cluster being closer to the STU population than to the other ITU subgroup. The PCA also shows a sibling pair with one sibling in the ITU population and the other in the STU population.

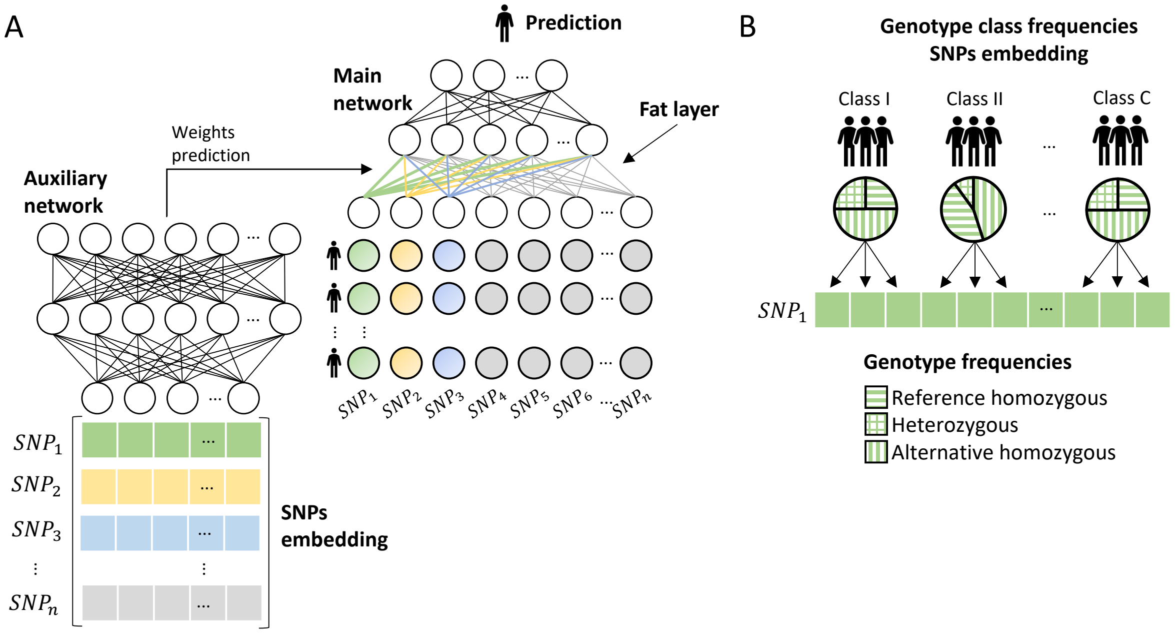


**Supplementary Figure 3: Diet Network architecture.** **A)** The Diet Network is composed of two neural networks trained jointly: a main network and an auxiliary network. The main network takes as input individual-level genotype data, encoded additively as 0, 1, or 2, representing the number of alternative alleles relative to the reference genome. The final output of the main network is a 24-dimensional vector corresponding to the 24 populations in the 1KGP (see Supplementary Results **S2.1** for description of the 24 populations), obtained after a softmax transformation to produce the Diet Network scores. The predicted population corresponds to the label with the highest DN score. Due to the high dimensionality of SNP data, this input generates a large number of weights in the initial layer, referred to as the "fat" layer. To reduce the number of trainable parameters and mitigate overfitting, the auxiliary network learns to predict the weights of this fat layer for each SNP. The auxiliary network receives as input a representation of each SNP, called the *SNPs embedding*. **B)** Example of the input provided to the auxiliary network for a single SNP ($SNP_1$). In our population classification task, SNP embeddings are computed from the genotype frequencies across populations. The input vector is constructed by concatenating the frequencies of each genotype (0, 1, 2) across the 24 populations from the 1KGP, resulting in a 72-dimensional vector ($24 \times 3 = 72$). Genotype frequencies are calculated using the training set within each cross-validation fold.

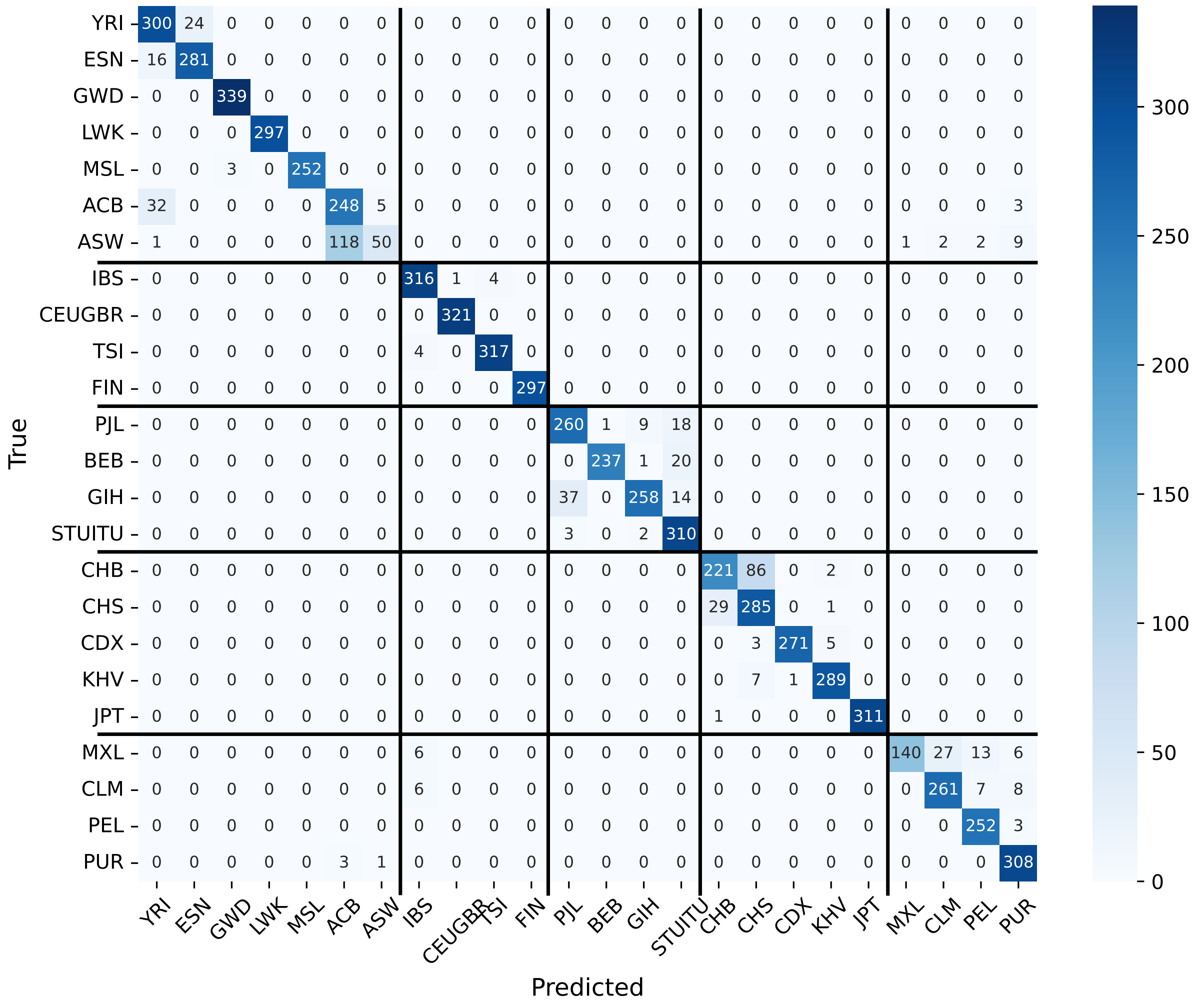


**Supplementary Figure 4: Diet Network models' predictions in 1KGP test sets.** Population labels predicted by fifteen Diet Networks models trained using 5-folds cross-validation across three random seeds, with an input dropout rate of 0.995. Black lines outline 1KGP super-population. The confusion matrix displays predictions on the held-out test set from each fold. As each individual appears once in the test set per cross-validation run and the procedure is repeated across three random seeds, each individual contributes three predictions to the confusion matrix.

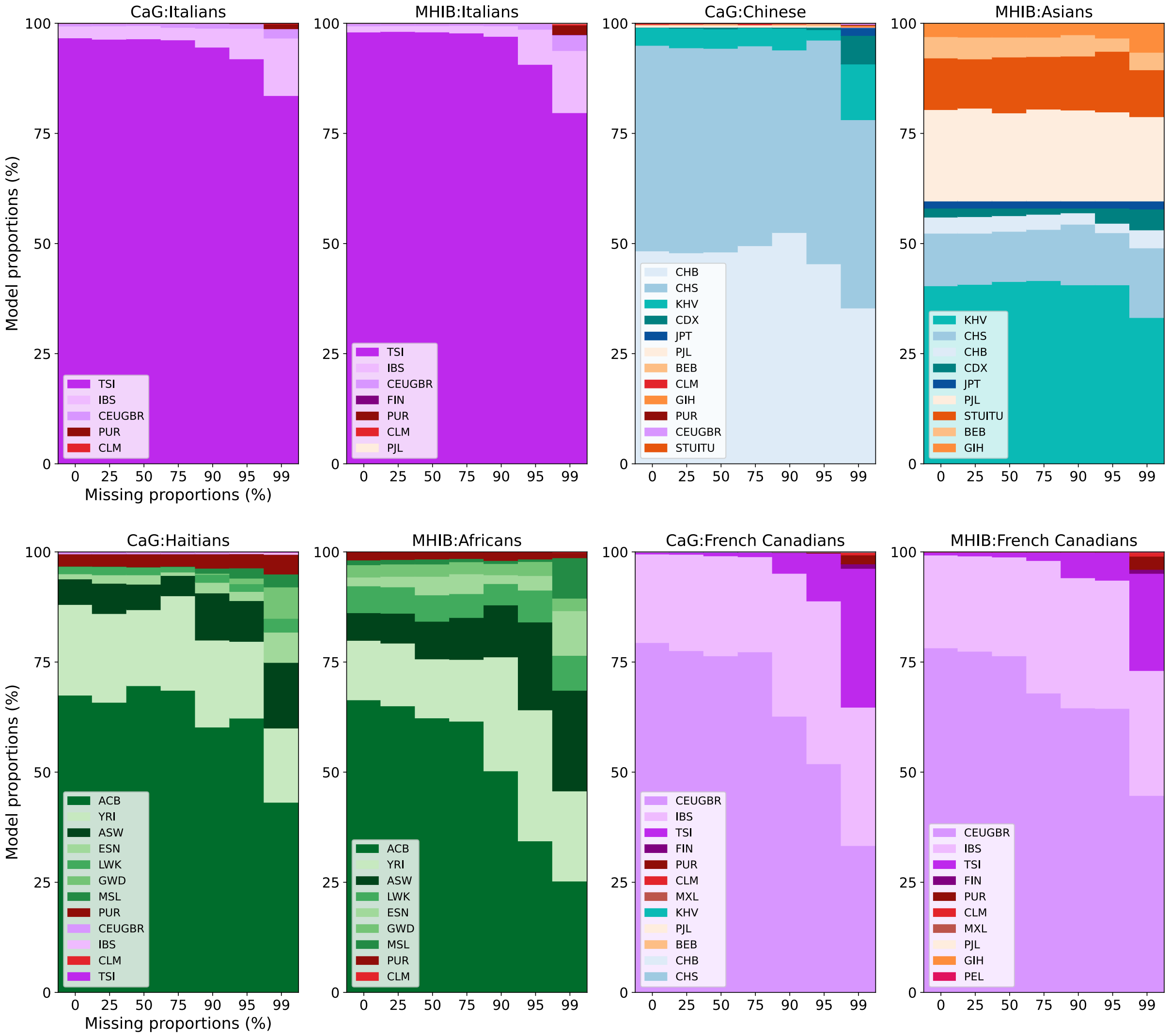


**Supplementary Figure 5: Generalizability in CaG and MHIB of Diet Network models trained with 0.995 input dropout across proportions of missing SNPs.** 1KGP predicted populations for each of selected proxy populations in CaG and MHIB. Each column represent the percentage of SNPs removed in the CaG and MHIB datasets to simulate missing values.

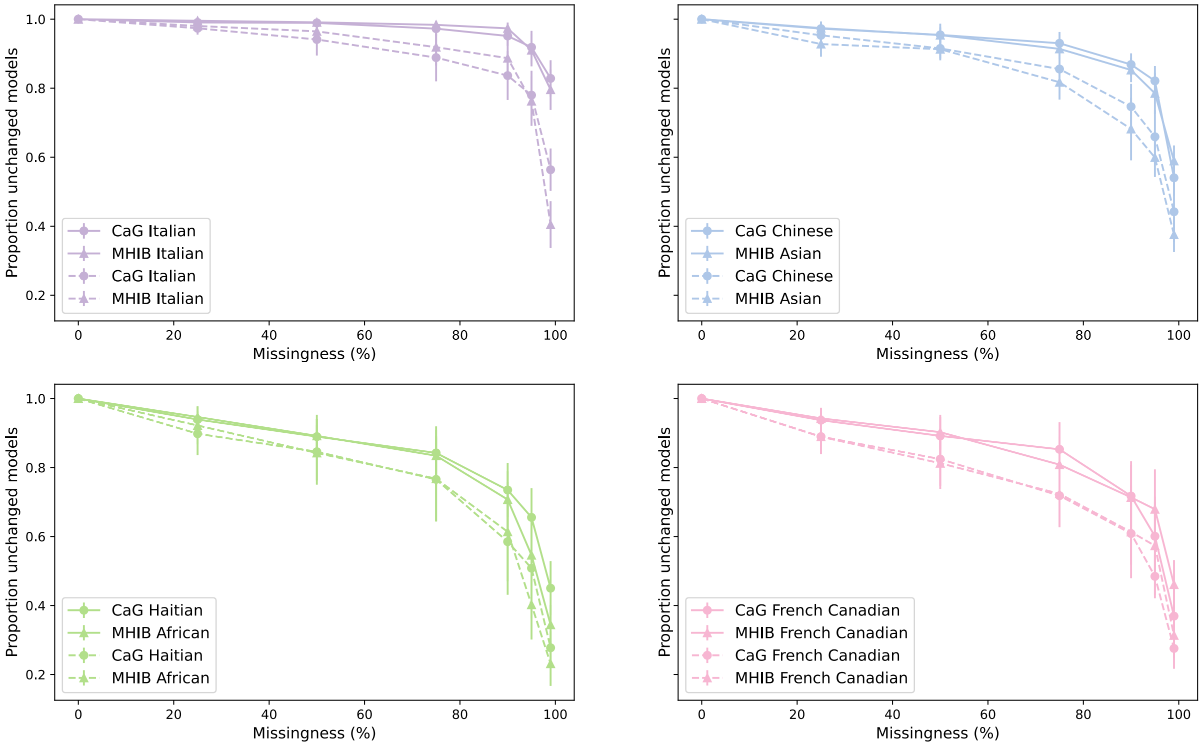


**Supplementary Figure 6: Input dropout impact on prediction changed across missing rates.** Proportion of models with a changed prediction across missing rates compared to predictions of models tested on complete SNPs data. Dashed lines: No input dropout. Plain lines : Input dropout = 0.995.

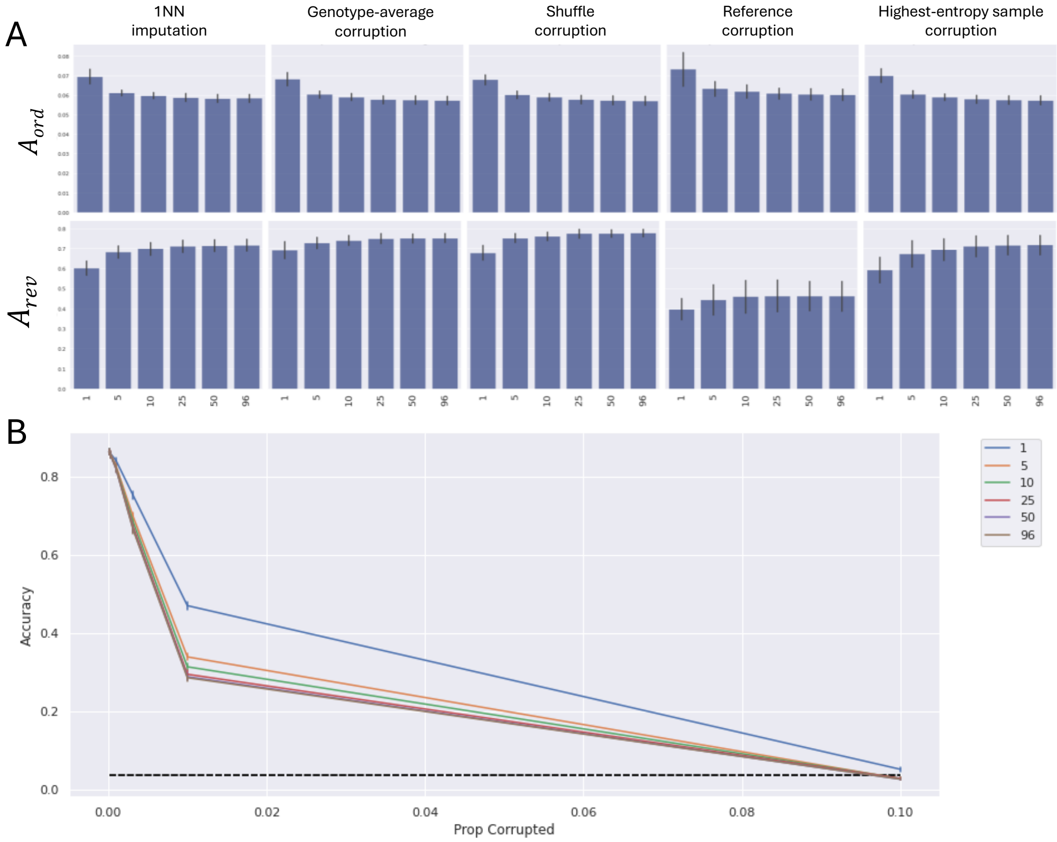


**Supplementary Figure 7: Area metrics for suboptimal baselines**. Suboptimal baseline sets were generated by sampling subsets of 1, 5, 10, 25, and 50 individuals from the full 96-sample multi-sample baseline. Attribution scores were computed using Integrated Gradients on fifteen Diet Network models trained via 5-fold cross-validation across three random seeds with 99.25% input dropout rate. (A) shows $A_{ord}$ and $A_{rev}$ metrics, where the single-sample baseline consistently performs worst. (B) shows classification accuracy for proportion of corrupted SNPs. A faster decline in accuracy is observed as the number of random samples included in the baseline set increases, reflecting the poorer ability of suboptimal baselines to identify important SNPs. Legend indicates the number of random samples in each baseline set. We show only up to 10% corruption for visual clarity.

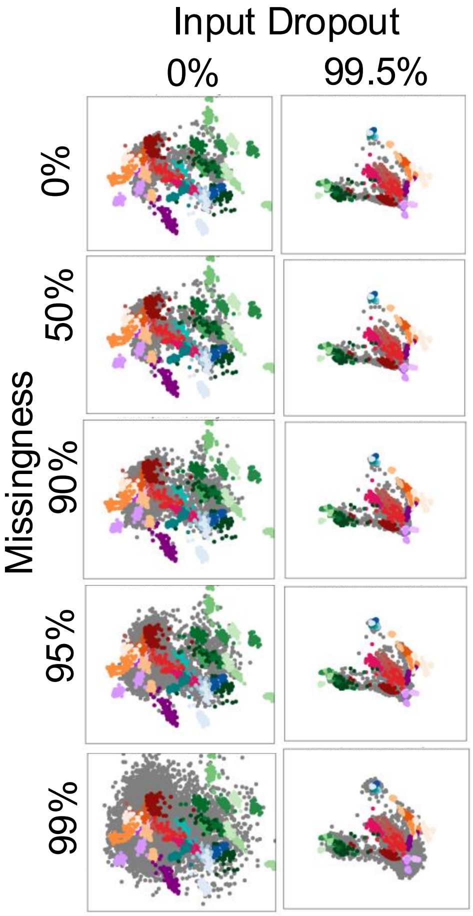


**Supplementary Figure 8: Diet Network model trained with input dropout yields more consistent hidden layer representations more consistent across different levels of missing SNPs.** PCA of values in the last hidden layer (100 neurons) of Diet Network models trained on 1KGP. The PCA was computed using 1KGP individuals from the test set and used to project the last hidden layer representations of all 17K individuals in the CaG biobank dataset. The plots show results for models trained without input dropout (left) and with an input dropout rate of 0.995 (right). Each row represents a different level of missingness, corresponding to the number of SNPs removed in the CaG dataset.

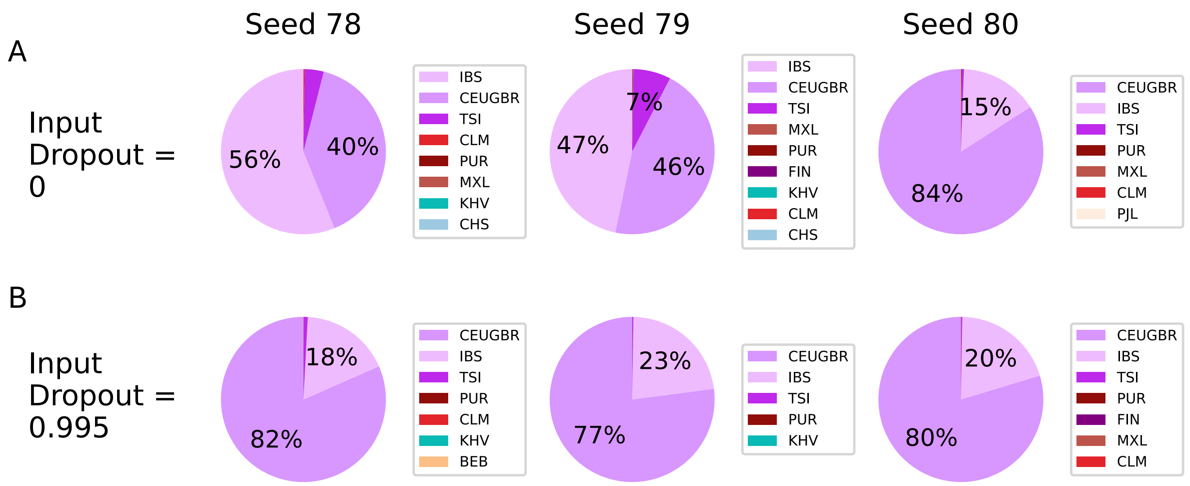


**Supplementary Figure 9: Diet Network models trained with input dropout yield more consistent predictions across random seeds** Diet Networks models predictions for CaG French Canadian individuals across three random seeds. Each pie chart represent predictions of 5 models (from the 5-folds cross-validation) for a given seed. (A) Predictions of Diet Network models trained without input dropout (B) Predictions of Diet Network models trained with a 0.995 input dropout rate.

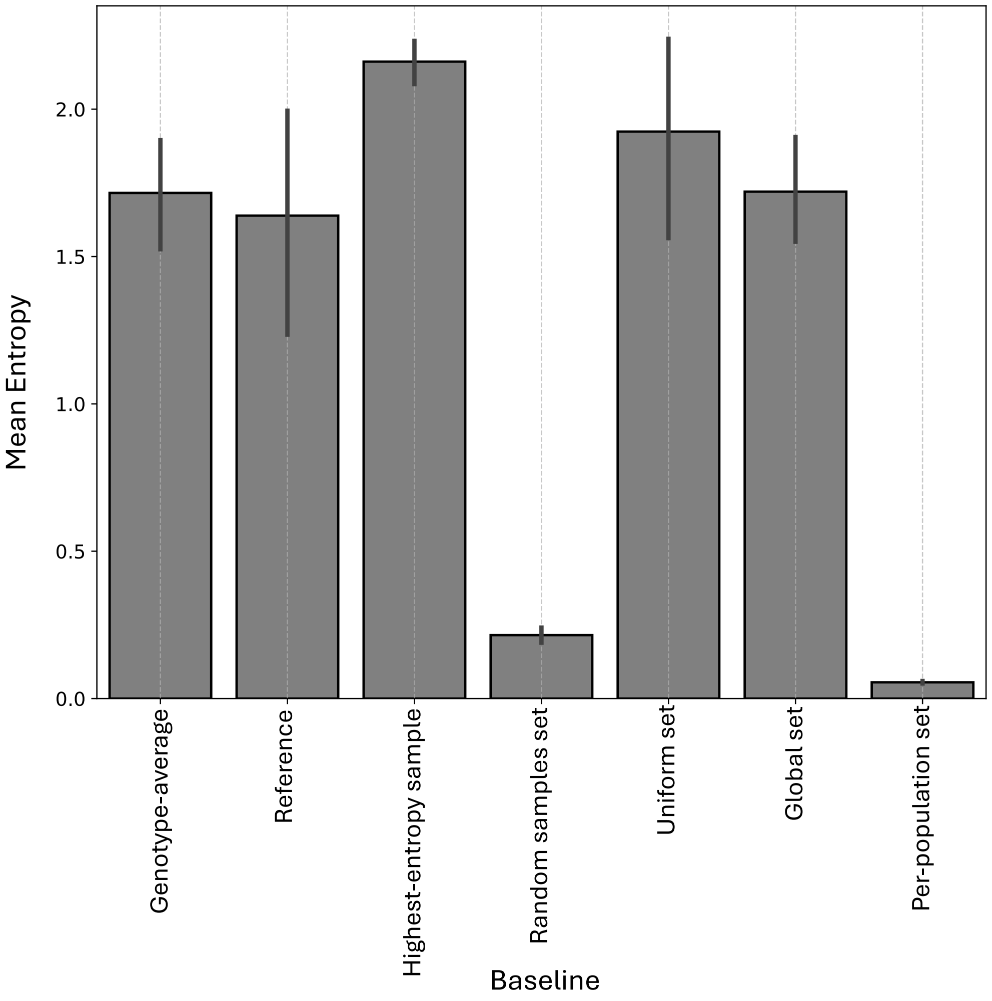


**Supplementary Figure 10: Prediction Entropy of Baselines**. Prediction entropy (y-axis) computed for each baseline input (x-axis) using fifteen models trained with 99.5% input dropout. For multi-sample sets, prediction entropy for all baselines in the set were counted.

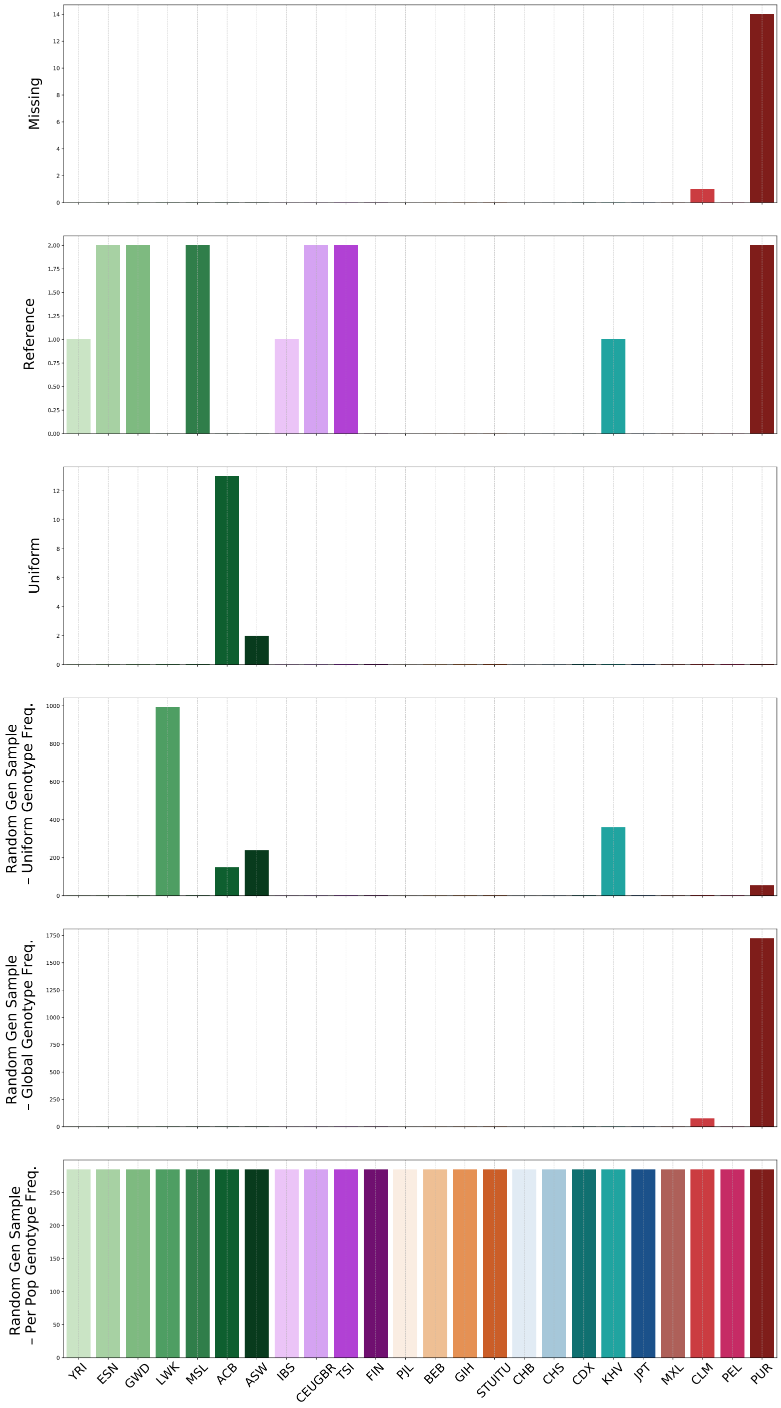


**Supplementary Figure 11: Prediction Diversity per Baseline**. Number of predicted classes (1KGP populations) for each baseline by the fifteen models trained with 99.5% input dropout. For multi-sample sets, we count prediction entropy for all baselines in the set ($15 models \times 96 baselines$ per set).

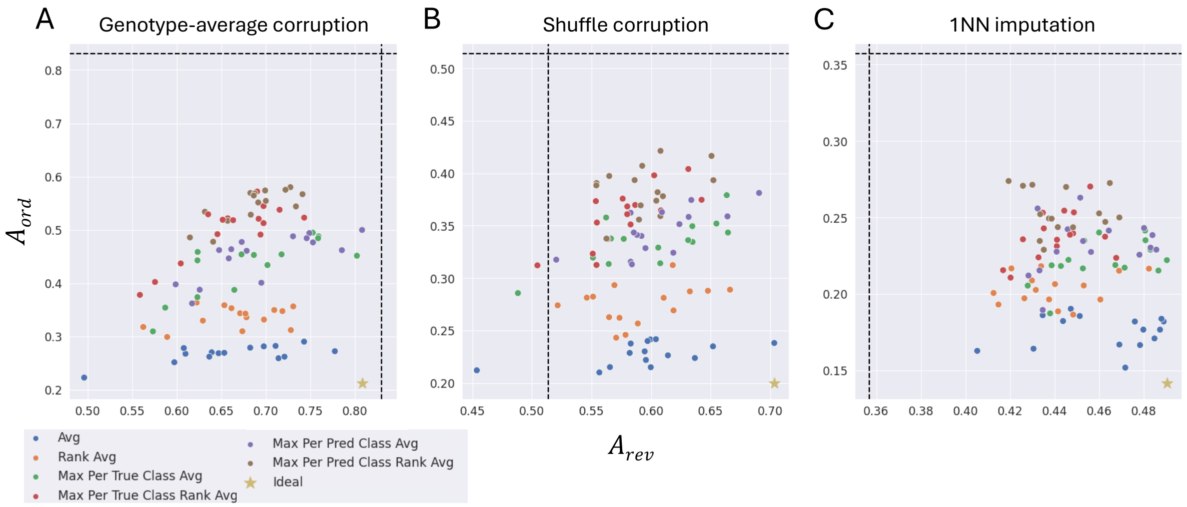


**Supplementary Figure 12: Global score aggregation strategies performance**. $A_{ord}$ (y-axis) and $A_{rev}$ (x-axis) computed for SNPs exclusion experiments performed using global scores computed for different local scores aggregation strategies. Local attribution scores were computed using Integrated Gradients and the genotype-average baseline on fifteen models trained with 99.25% input dropout. We compare the models' area metrics against an ideal score (designated by a star in the bottom right corner). Horizontal and vertical lines represent $A_{ran}$, the area under the curve obtained when corrupting SNPs at random. SNPs were corrupted using (A) genotype-average corruption, (B) shuffle corruption and (C) 1NN imputation.

### S4 Supplementary Tables

Supplementary Table 1: Thousand Genomes (1KG) Project Populations Code and Description. Officially approved population descriptors were established to accurately represent both the ancestral geography or ethnicity of each population and the geographic location where the samples were collected. These descriptors incorporate input from sample donor communities regarding how they wish to be described and should be used as designated without modification.

| **Code** | **Superpopulation code** | **Description** |
|---|---|---|
| YRI | AFR | Yoruba in Ibadan, Nigeria |
| ESN | AFR | Esan in Nigeria |
| GWD | AFR | Gambian in Western Division, The Gambia - Mandinka |
| LWK | AFR | Luhya in Webuye, Kenya |
| MSL | AFR | Mende in Sierra Leone |
| ACB | AFR | African Caribbean in Barbados |
| ASW | AFR | African Ancestry in Southwest US |
| IBS | EUR | Iberian populations in Spain |
| CEU | EUR | Utah residents (CEPH) with Northern and Western European ancestry |
| GBR | EUR | British in England and Scotland |
| TSI | EUR | Toscani in Italy |
| FIN | EUR | Finnish in Finland |
| PJL | SAS | Punjabi in Lahore, Pakistan |
| BEB | SAS | Bengali in Bangladesh |
| GIH | SAS | Gujarati Indians in Houston, TX |
| STU | SAS | Sri Lankan Tamil in the UK |
| ITU | SAS | Indian Telugu in the UK |
| CHB | EAS | Han Chinese in Beijing, China |
| CHS | EAS | Han Chinese South |
| CDX | EAS | Chinese Dai in Xishuangbanna, China |
| KHV | EAS | Kinh in Ho Chi Minh City, Vietnam |
| JPT | EAS | Japanese in Tokyo, Japan |
| MXL | AMR | Mexican Ancestry in Los Angeles, California |
| CLM | AMR | Colombian in Medellin, Colombia |
| PEL | AMR | Peruvian in Lima, Peru |
| PUR | AMR | Puerto Rican in Puerto Rico |

Codes and descriptions are provided by the International Genome Sample Resource [8] and listed in the order used in the paper figures. Superpopulation codes: African ancestry (AFR), Americas (AMR), East Asian ancestry (EAS), European ancestry (EUR), South Asian ancestry (SAS)

Supplementary Table 2: Area metrics across baselines for corruption methods that replace corrupted SNPs with corresponding values found in single-sample baselines

| | **Reference Corruption** | | **Genotype-average Corruption** | | **Highest-entropy sample Corruption** | |
|---|---|---|---|---|---|---|
| **Baseline** | $A_{ord}$ | $A_{rev}$ | $A_{ord}$ | $A_{rev}$ | $A_{ord}$ | $A_{rev}$ |
| Reference | 0.02 ± 0.01 | 0.85 ± 0.03 | 0.05 ± 0.01 | 0.92 ± 0.01 | 0.04 ± 0.01 | 0.79 ± 0.04 |
| Genotype-average | 0.03 ± 0.01 | 0.8 ± 0.05 | 0.04 ± 0.01 | 0.92 ± 0.01 | 0.04 ± 0.01 | 0.87 ± 0.02 |
| Highest-entropy sample | 0.04 ± 0.01 | 0.77 ± 0.07 | 0.05 ± 0.01 | 0.92 ± 0.01 | 0.03 ± 0.01 | 0.88 ± 0.01 |
| Set - Random samples | 0.03 ± 0.01 | 0.8 ± 0.05 | 0.04 ± 0.01 | 0.92 ± 0.01 | 0.04 ± 0.01 | 0.87 ± 0.02 |
| Set - Uniform frequency | 0.06 ± 0.01 | 0.73 ± 0.06 | 0.05 ± 0.01 | 0.92 ± 0.01 | 0.05 ± 0.01 | 0.82 ± 0.03 |
| Set - Global frequency | 0.03 ± 0.01 | 0.8 ± 0.05 | 0.04 ± 0.01 | 0.92 ± 0.01 | 0.04 ± 0.01 | 0.87 ± 0.02 |
| Set - Per-population frequency | 0.03 ± 0.01 | 0.8 ± 0.05 | 0.04 ± 0.01 | 0.92 ± 0.01 | 0.04 ± 0.01 | 0.87 ± 0.02 |

Supplementary Table 3: Area metrics for corruption order based on SNPs global attribution scores.

| Corruption Style | Score | $A_{ord}$ | $A_{rev}$ |
|---|---|---|---|
| Genotype-average | Avg | $0.2679 \pm 0.0160$ | $0.6621 \pm 0.0703$ |
| | Rank Avg | $0.3368 \pm 0.0196$ | $0.6667 \pm 0.0494$ |
| | Max Per True Class Avg | $0.4329 \pm 0.0530$ | $0.6821 \pm 0.0680$ |
| | Max Per True Class Rank Avg | $0.5000 \pm 0.0536$ | $0.6612 \pm 0.0516$ |
| | Max Per Pred Class Avg | $0.4509 \pm 0.0429$ | $0.6950 \pm 0.0637$ |
| | Max Per Pred Class Rank Avg | $0.5460 \pm 0.0320$ | $0.6865 \pm 0.0375$ |
| | Random SNP Corruption | $0.8300 \pm 0.0174$ | |
| Shuffle | Avg | $0.2277 \pm 0.0111$ | $0.5958 \pm 0.0533$ |
| | Rank Avg | $0.2752 \pm 0.0188$ | $0.5905 \pm 0.0405$ |
| | Max Per True Class Avg | $0.3334 \pm 0.0226$ | $0.5973 \pm 0.0506$ |
| | Max Per True Class Rank Avg | $0.3586 \pm 0.0277$ | $0.5780 \pm 0.0345$ |
| | Max Per Pred Class Avg | $0.3449 \pm 0.0217$ | $0.6068 \pm 0.0400$ |
| | Max Per Pred Class Rank Avg | $0.3842 \pm 0.0229$ | $0.5955 \pm 0.0300$ |
| | Random SNP Corruption | $0.5138 \pm 0.0244$ | |
| 1NN imputation | Avg | $0.1752 \pm 0.0109$ | $0.4624 \pm 0.0255$ |
| | Rank Avg | $0.2033 \pm 0.0104$ | $0.4401 \pm 0.0197$ |
| | Max Per True Class Avg | $0.2215 \pm 0.0139$ | $0.4613 \pm 0.0202$ |
| | Max Per True Class Rank Avg | $0.2377 \pm 0.0158$ | $0.4414 \pm 0.0145$ |
| | Max Per Pred Class Avg | $0.2317 \pm 0.0179$ | $0.4567 \pm 0.0210$ |
| | Max Per Pred Class Rank Avg | $0.2540 \pm 0.0144$ | $0.4429 \pm 0.0151$ |
| | Random SNP Corruption | $0.3569 \pm 0.0082$ | |

[3] Adriana Romero, Pierre Luc Carrier, Akram Erraqabi, Tristan Sylvain, Alex Auvolat, Etienne Dejoie, Marc-Andre Legault, Marie-Pierre Dube, Julie Hussin, and Yoshua Bengio. Diet networks: Thin parameters for fat genomics. *ICLR*, 2017.

[4] Karen Simonyan, Andrea Vedaldi, and Andrew Zisserman. Deep inside convolutional networks: Visualising image classification models and saliency maps. In Yoshua Bengio and Yann LeCun, editors, *2nd International Conference on Learning Representations, ICLR 2014, Banff, AB, Canada, April 14-16, 2014, Workshop Track Proceedings*, 2014.

[5] Héléna A. Gaspar and Gerome Breen. Probabilistic ancestry maps: a method to assess and visualize population substructures in genetics. *BMC Bioinformatics*, 20(1), March 2019.

[6] Dhriti Sengupta, Ananyo Choudhury, Analabha Basu, and Michèle Ramsay. Population stratification and underrepresentation of indian subcontinent genetic diversity in the 1000 genomes project dataset. *Genome Biology and Evolution*, 8(11):3460–3470, October 2016.

[7] Daniel Schlauch, Heide Fier, and Christoph Lange. Identification of genetic outliers due to sub-structure and cryptic relationships. *Bioinformatics*, 33(13):1972–1979, March 2017.

[8] Susan Fairley, Ernesto Lowy-Gallego, Emily Perry, and Paul Flicek. The international genome sample resource (igsr) collection of open human genomic variation resources. *Nucleic Acids Research*, 48(D1):D941–D947, October 2019.
